# A high throughput bispecific antibody discovery pipeline

**DOI:** 10.1101/2021.09.07.459213

**Authors:** Aude I. Segaliny, Jayapriya Jayaraman, Xiaoming Chen, Jonathan Chong, Ryan Luxon, Audrey Fung, Qiwei Fu, Xianzhi Jiang, Rodrigo Rivera, Xiaoya Ma, Ci Ren, Jan Zimak, Per Niklas Hedde, Yonglei Shang, George Wu, Weian Zhao

**Author notes:** co-first authors (Aude I. Segaliny, Jayapriya Jayaraman). co-second authors (Xiaoming Chen, Jonathan Chong, Ryan Luxon). **Corresponding authors:** George Guikai Wu, Weian Zhao.

## Abstract

Bispecific antibodies (BsAbs) represent an emerging class of immunotherapy but inefficiency in the current BsAb discovery paradigm has limited their broad clinical availability. Here we report a high throughput, agnostic, single-cell-based BsAb functional screening pipeline, comprising molecular and cell engineering for efficient generation of BsAb library cells, followed by functional interrogation at the single-cell level to identify and sort positive clones and downstream sequence identification with single-cell PCR and sequencing and functionality characterization. Using a CD19xCD3 bispecific T cell engager (BiTE) as a model system, we demonstrate that our single cell platform possesses a high throughput screening efficiency of up to one and half million variant library cells per run and can isolate rare functional clones at low abundance of 0.008%. Using a complex CD19xCD3 BiTE-expressing cell library with approximately 22,300 unique variants comprising combinatorially varied scFvs, connecting linkers and VL/VH orientations, we have identified 98 unique clones including extremely rare ones (∼ 0.001% abundance). We also discovered BiTEs that exhibit novel properties contradictory to conventional wisdom, including harboring rigid scFv connecting peptide linkers yet with in vitro cytotoxicity comparable to that of clinically approved Blinatumomab. Through sequencing analyses on sorted BiTE clones, we discovered multiple design variable preferences for functionality including the CD19_VL-VH_– CD3_VH-VL_ and CD19_VH-VL_–CD3_VH-VL_ arrangements being the most favored orientation. Sequence analysis further interrogated the sequence composition of the CDRH3 domain in scFvs and identified amino acid residues conserved for function. We expect our single cell platform to not only significantly increase the development speed of high quality of new BsAb therapeutics for cancer and other disorders, but also enable identifying generalizable design principles for new BsAbs and other immunotherapeutics based on an in-depth understanding of the inter-relationships between sequence, structure, and function.

## Introduction

Bispecific antibodies (BsAbs) represent a highly attractive class of antibodies and immunotherapeutics that hold great potential to treat many disorders including cancer and autoimmunity^1–5^. They are unnatural biologics that are engineered to recognize two different epitopes either on the same or different target antigens. BsAbs offer unique therapeutic modes of action such as activating and engaging immune cells to kill tumor cells and simultaneously acting on two synergizing therapeutic targets^5–8^. Therefore, BsAbs can exhibit superior therapeutic advantages over traditional monoclonal antibodies (mAbs) with respect to specificity, efficacy, toxicity and drug resistance. T cell activating BsAbs (TABs) represent the largest subclass of BsAbs and account for approximately 80% of the BsAb preclinical and clinical pipeline^9^. Indeed, among the three approved BsAb drugs, two are TABs: 1) Blinatumomab, targeting CD3*ε* and tumor antigen CD19, and 2) Catumaxomab, targeting T cell receptor (TCR) subunit CD3*ε* and tumor antigen EpCAM^9,10^. TABs can directly link T cells and tumor cells for killing and hold great potential to treat many types of cancer. There are currently over 45 CD3-based TABs including BCMAxCD3, Her2xCD3, CEAxCD3, and PSMAxCD3 being clinically tested for treatment of solid and hematological cancers^1,6^. TABs can be designed in several formats including bispecific T cell engagers (BiTEs) which use tandemly linked single-chain fragment variable fragments (scFvs) targeting a T cell epitope and tumor antigen and common-light-chain IgG with a full-size immunoglobulin configuration where each variable heavy chain targets either a T-cell receptor or a tumor antigen^1,6^.

Unfortunately, developing therapeutic BsAbs, including TABs remains to be an extremely challenging, slow and costly process, limiting their clinical availability for a broad range of diseases. We attribute this bottleneck largely to the complexity and inefficiency of the current BsAb discovery methods. The conventional workflow for BsAb discovery commonly starts with characterizing a pool of mAb binders for two respective antigens or epitopes. Then a panel of mAbs are selected for engineering BsAb variants individually by joining two antigen-binding domains derived from each parental mAb via subunit heterodimerization or direct genetic fusion (*e*.*g*., BiTEs)^11–23^. Then the individual BsAb variants are expressed, purified, and subsequently tested individually in a dual-target binding assay and/or a functional assay to identify positive clones. Of note, despite that BsAb construction usually starts with two pre-characterized antigen binders, integrating them to become a functional BsAb is much less straightforward. Indeed, the binding activity of a BsAb to its targets is only a prerequisite, which does not always translate into a good functional activity. For TABs, specifically, the T cell activation and tumor-killing functions of resulting BsAbs depend crucially on numerous factors including epitope location and distance to cell membrane^24–29,^ antigen size and density^27–36^, antibody/antigen binding affinity^31–35,37,38^ the size, valency, folding, structural orientation and mechanics of BsAbs, and the linker length and flexibility^27,36,39–42^. While it is well recognized that small variations in BsAbs’ formats and characteristics can be critical determinants of functionality, the relationships among these factors are complicated, and currently there is no general design principle that can inform BsAb function. Practically, therefore, the field has been relying on a trial-and-error process in which BsAbs are empirically designed and constructed from existing mAb binders and must then be evaluated individually for their functionality. This conventional approach, typically involving a microtiter-plate and liquid handling system, is biased and significantly limits BsAb discovery throughput, success rate and turnaround time to identify a therapeutic lead for further development.

In this work, we describe a high throughput, single-cell-based BsAb functional screening pipeline (Fig. 1) that can directly interrogate the functions of a large number of individual BsAb variants from an unbiased library. The single cell pipeline integrates several modules, including a molecular and cell engineering module for efficient generation of BsAb-variant cell libraries and reporter cells, an opto-electro-mechanical module for droplet based single-cell functional screening to identify and sort desired BsAb clones, and a downstream molecular analysis and validation module to determine the sequences and functionality of the candidate BsAbs (Fig. 1). At its core, the single cell platform employs a droplet microfluidic-based system to compartmentalize and interrogate individual BsAb-producing cells with co-encapsulated cell reporters. Functional BsAb clones will be able to activate cell reporters to produce fluorescence, which allows “positive” droplets to be detected and sorted from a heterogeneous population. Of note, we have implemented innovative multiplexed orthogonal assay chemistry, and multi-point detection and droplet-indexing strategy to ensure screening fidelity. Using an established CD19xCD3 BiTE as a model system, we describe below how each module is constructed in a streamlined workflow. We have demonstrated that our single cell platform can accommodate one and half million library cells in a single run, which is 2-3 orders of magnitude higher in throughput than that of conventional methods (10s-100s of clones for manual microtiter-plate handling methods or 1,000s-10,000s of clones for robotics-liquid handling systems). Using a spiked library prepared with HEK293 cells that expresses human CD19 at a comparable level to lymphoma cell lines, and secretes a functional CD19xCD3 BiTE, we characterized the analytical performance including screening efficiency and demonstrated the single cell platform is capable of isolating ≥ 1 copies of a rare functional clone (0.008% abundance) with 95% confidence in a single run. Using a complex CD19xCD3 BiTE library with approximately 22,300 unique variants where we combinatorially varied scFv binders encompassing different epitopes and affinities, VL/VH orientations, and length and flexibility of scFv connecting linkers, we have identified 98 unique functional clones including extremely rare clones (∼ 0.001% abundance) and clones bearing rigid scFv connecting peptide linkers that are against conventional wisdom. Sequence analysis of sorted clones further interrogated preferences of different design variables for functionality and discovered that the CD19_VL-VH_–CD3_VH-VL_ and CD19_VH-VL_–CD3_VH-VL_ arrangements are the most favored orientation. Sequence analysis further revealed the sequence composition of the CDRH3 domain at single residue resolution and identified amino acid residues conserved or promiscuous for function.

**Figure 1.**
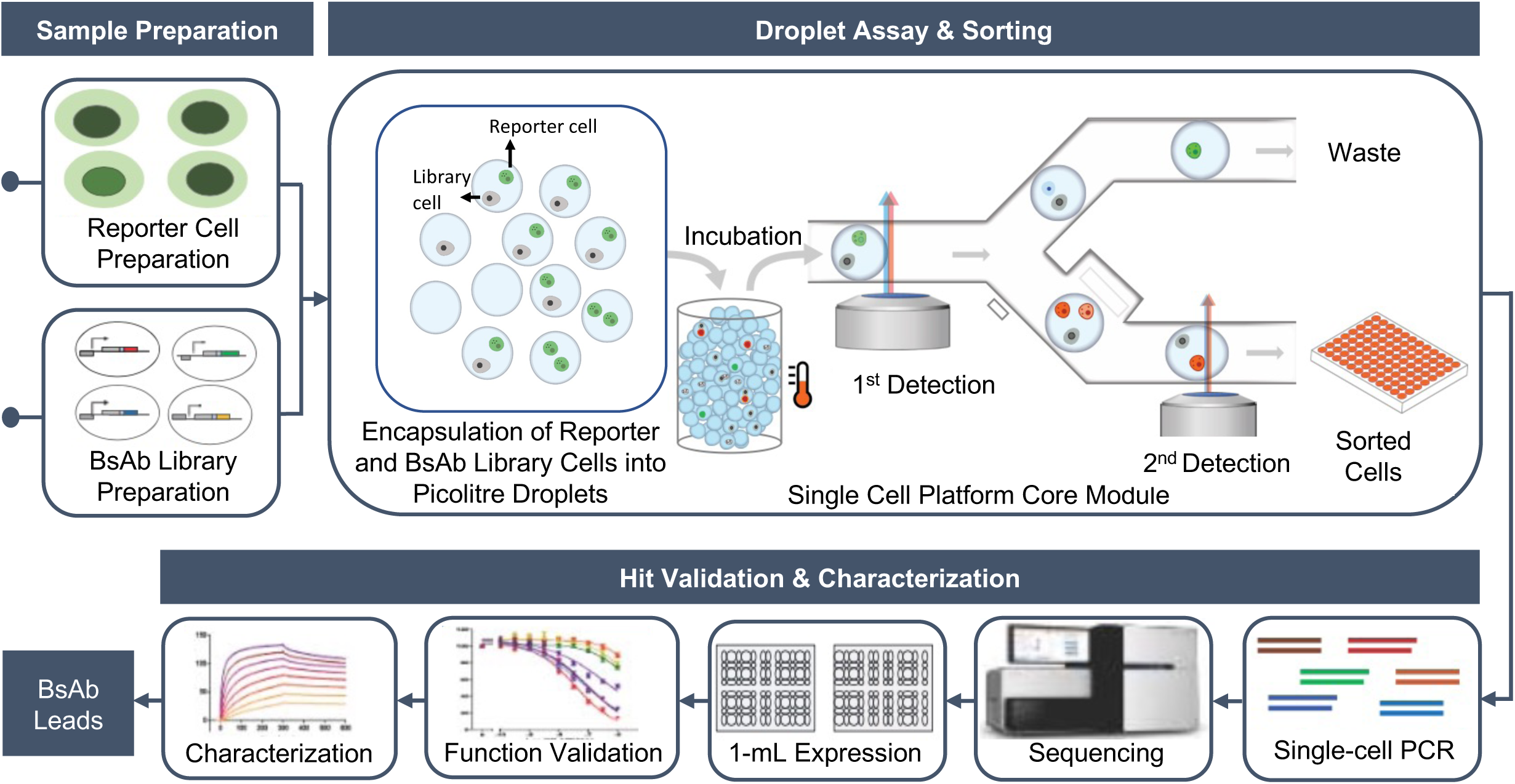
Overview of single-cell-based BsAb discovery pipeline.

## Results

### BsAb structural and compositional considerations and combinatorial library design

The structural and compositional attributes of BsAb design have a significant impact on their functionalities. Unlike the trial-and-error approach used in conventional methods, we can conduct unbiased interrogation of many structural and compositional variables (*e*.*g*., affinity, epitope location, linker design and relative orientations of the binding antibody moieties) combinatorially without prior knowledge owing to our single cell platform’s high throughput. In this study, we sought to develop and validate the platform using CD19xCD3 BiTE as a model because it has been clinically validated and widely studied (*i*.*e*., Blinatumomab), before we expand its utility to other BsAbs in the future. BiTEs consist of tandem single-chain fragment variables (scFvs) with one scFv binding to a T cell receptor subunit (most commonly CD3*ε*), and the other to a target antigen (*e*.*g*., a tumor associated antigen) (Fig. 2a). The two scFvs are linked by a peptide linker. Each scFv comprises a VH and a VL domain derived from mAb binders, which are linked by a short peptide linker. Engaging T cells with tumor cells in close proximity using BiTEs activates T cells to release cytokines and apoptotic factors (mainly perforins and granzymes) that subsequently can kill tumor cells.

**Figure 2.**
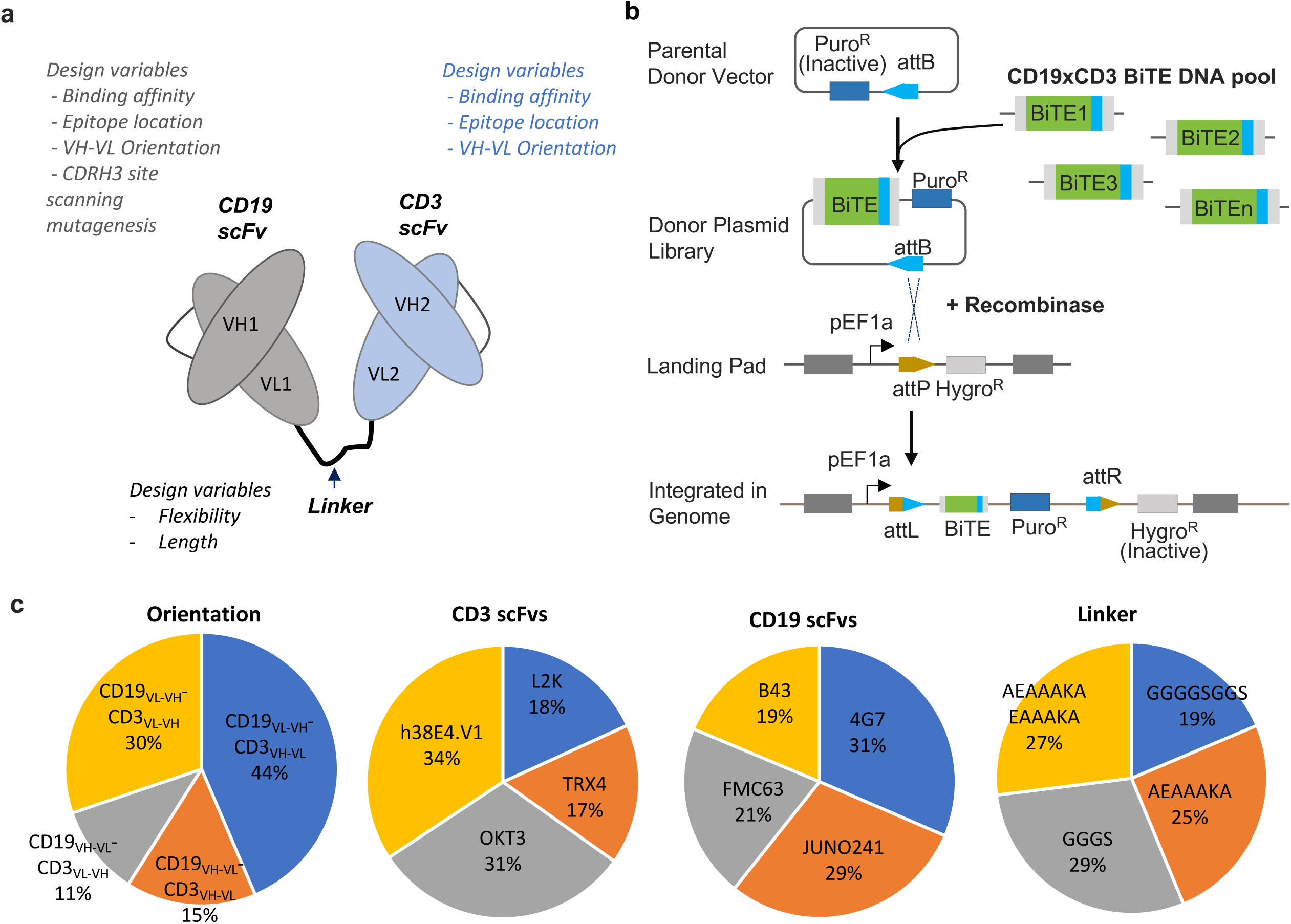
BiTE library generation and characterization of integrated library diversity. **(a)** Design variables interrogated for the CD19 scFv ,CD3 scFv and peptide linker connecting the scFvs. **(b)** Schematic representation of integration of single copy BiTE variant in landing pad harboring HEK293 cell line (HEK293_LP) with promoterless BiTE gene encoding donor plasmid library. **(c)** Representation of diversity of BiTE design features obtained from long read amplicon sequencing (PacBio SMRT™)of PCR amplified BiTE genes from genomic DNA of BiTE integrated HEK293_LP cells, (*from left to right*) Distribution of relative orientation of scFvs, Distribution of relative CD3 scFvs, Distribution of CD19 scFvs and Distribution of scFv interconnecting linkers. ∼22,300 unique BiTE sequences obtained from long read amplicon sequencing was used for analyzing distribution of design parameters (relative scFv orientation, CD3scFv type, CD19 scFv type and scFv interconnecting linker type). Genomic DNA template used for amplifying integrated BiTE genes was obtained from 3×10^6^ HEK_LP^CD19/BiTE^ cells obtained from *n=1* integration experiment.

Our library design incorporates multiple variables including choice of scFv targeting various epitopes, scFv binding affinities, length and flexibility of linkers, and orientation of scFvs, all known determinants of BiTE functionality (Fig. 2a). For CD19 binders we chose scFv sequences based on mAb 4G7, mAb FMC63, mAb B43 and mAb Juno241 (listed Table 1 with their characteristics). mAb B43 is the basis of Blinatumomab^43,44^. mAb 4G7 and mAb FMC63 bind to a conformationally similar epitope on hCD19, but distinct from that of B43 binding epitope^45^. While 4G7 and B43 have reportedly similar affinities, FMC63 has at least one order of magnitude higher affinity^43,46–48^. Juno241 has been shown to have strong binding to CD19^49^. We also similarly adapted four CD3 mAb binders OKT3, L2K, h38E4.v1 and TRX4 (listed in Table 2 with their characteristics). L2K, for instance shares over 90% sequence identity with OKT3 and recognizes the same epitope on CD3*ε* but differs in affinity by about 100-fold^50,51^. L2K is the basis of clinically approved Blintumomab molecule^43^, while OKT3 and TRX4 are commercially available as mAb formats treatment of certain auto-immune conditions^52^. TRX4 and h38E4.v1 have a higher affinity to CD3 compared to OKT3 and L2K^43,51,53,54^. To increase library diversity, we performed a site scanning mutagenesis with NNW degenerate codons on the CDRH3 region of the CD19 scFv candidates (see Materials and Methods). For peptide linkers that connect the two scFvs, we chose flexible short and long linkers based on GGGGS motif as well as rigid short and long linkers based on EAAAK motif (Table 3)^55,56^. The theoretical diversity of our combinatorial library, calculated as the product of number of distinct CD19 and CD3 variants, number of scFv interconnecting linkers, and number of different VL/VH orientations is 47,880 (Table 4). A mammalian cell-based BiTE expression system incorporating this diverse library is then constructed (below) to prepare input BsAb-variant expressing cell library for the single cell screening pipeline.

**Table 1.** aCD19 scFvs and their characteristics such as affinity, epitope and CDRH3 lengths. The diversity generated in CD19 scFvs by performing site scanning mutagenesis on CDRH3 residues and altering relative orientation of VH-VL regions is also shown.

| aCD19 scFv | Affinity (nM) | Epitope of binding | CDRH3 length | CDRH3-NNW randomized sequences/scFv | VH/VL orientation |
| --- | --- | --- | --- | --- | --- |
| 4G7 | 8.4 nM | Conformationally similar epitope with FMC63, partially overlapping yet distinct from B43. | 12 | 216 | 2 |
| FMC63 | 0.4 nM | Discontinuous epitope partially overlapping with B43. | 12 | 216 | 2 |
| Juno 241 | NA | NA | 16 | 288 | 2 |
| B43 | 1.5 nM | Loop 216-224, Loop 155-166 on CD19 | 15 | 270 | 1 |
| Total Number of aCD19 variants |  |  |  |  | 1710 |

**Table 2.** aCD3 scFvs and their characteristics such as affinity, epitope of binding. Diversity in aCD3 scFvs was generated by altering relative orientation of VH-VL regions.

| aCD3 scFv | Affinity (nM) | Epitope of binding | CDRH3/Rand omization | VH/VL orientation |
| --- | --- | --- | --- | --- |
| OKT3 | 0.5 nM | Discontinuous epitope on CD3ε, middle of extracellular domain near a charged patch | No randomization of CDRH3 residues | 2 |
| L2K | 110 nM | Overlapping epitope with OKT3 |  | 2 |
| TRX4 | 0.09 nM | NA |  | 2 |
| h38E4.v1 | 1 nM | N-terminal portion of CD3ε extracellular domain |  | 1 |
| Total Number of aCD3 variants |  |  |  | 7 |

**Table 3.**
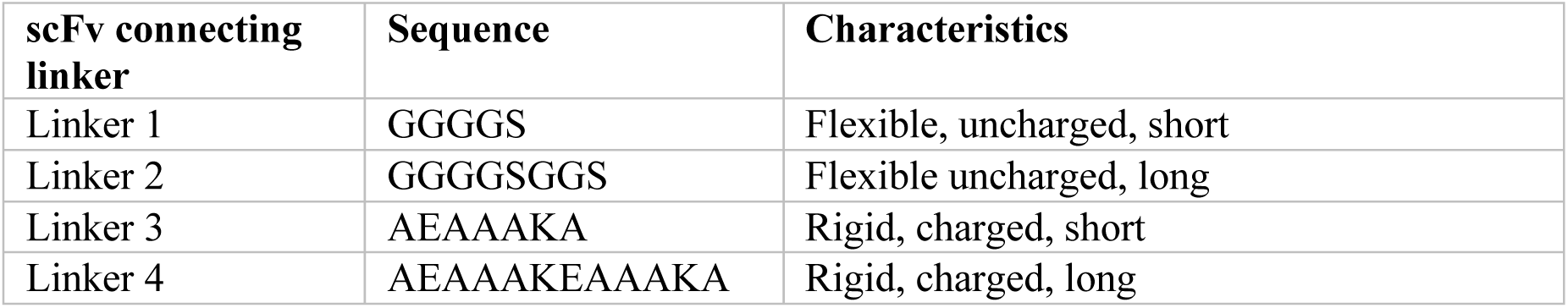
scFv interconnecting linkers chosen for library generation. Linkers were varied based on flexibility, charge and length.

**Table 4.**
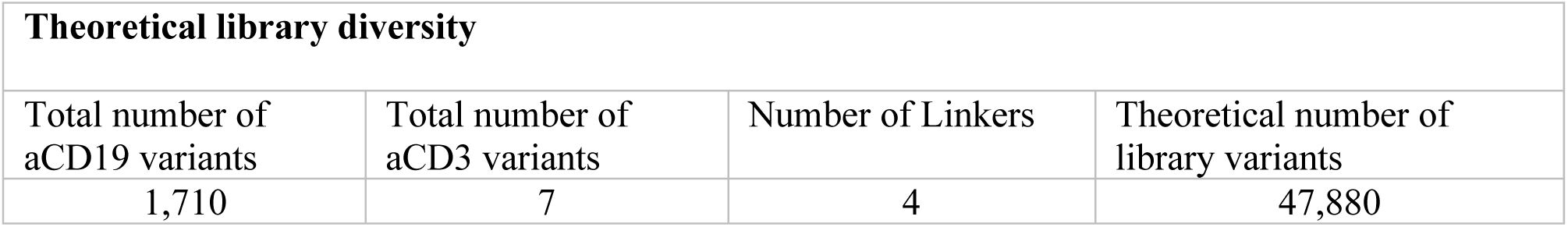
Theoretical library diversity of BiTE library calculated as product of number of aCD19 variants, number of aCD3 variants and number of linkers.

### Construct mammalian cell-based BsAb-variant expressing library cells encoding a single copy each for a BsAb variant and expressing target antigens

We conceived a screening strategy where reporter T-cells will be encapsulated in droplets with a single BiTE secreting cell that also expresses the target antigen of interest, human CD19, on the surface at levels comparable to common lymphoma cell lines. Having CD19 and BiTE expressed in the same cell, rather than in two different cells, allows us to achieve adequate droplet co-encapsulation efficiency with the reporter cell due to Poisson distribution^57^. We expect that functional BiTEs secreted from the CD19 expressing cells will crosslink the cell surface CD19 and TCR-CD3 complex on the T cell reporter cells and produce an optically detectable intra-droplet reporter signal. To ensure that we can connect intra-droplet reporter signal (phenotype) to a specific BiTE sequence (genotype), it is crucial that only one type of BiTE variant is expressed from each CD19 expressing cell. To accomplish this, we sought to develop a mammalian (HEK293) cell-based BiTE variant library expression platform where each cell in the library is genetically integrated with landing-pad expression vehicle that encodes a single copy each for a BsAb variant. HEK293 was chosen here in part because of its ease of use for small-scale expression as well as scaled production for recombinant proteins including BsAbs. Specifically, we adopted a commercially available HEK293 landing-pad parental line (“HEK293_LP”) that has an integrated “landing-pad” component in the genome (*i*.*e*., single BxB1 recombinase-attP attachment site in Hipp11 (H11) safe-harbor locus on Chromosome 22 in the human genome)^58^. The compatibility of this HEK293 landing-pad line for efficient integration (up to 38%) of an exogeneous gene-of-interest is validated by mCherry gene insertion in the landing-pad locus (Supplementary Fig. 1a).

We next prepared a target cell line that expresses human CD19 and a functional CD19xCD3 BiTE. We first stably integrated a CD19 expression cassette in a safe harbor locus on Chromosome 1 using CRISPR-Cas9 in the HEK293_LP cell line^59^. We then, clonally selected CD19 engineered HEK293_LPs for CD19 expression levels comparable to other standard CD19+ cells including Raji, NALM-6, and SU-DHL-6 (Supplementary Fig. 1b). Lastly, we created a BiTE-expressing cell library with a size and diversity that recapitulate those of typical antibody germline alleles and epitopes for a given target. For this, we first generated a “donor” plasmid library pool which harbored combinatorially combined BiTE variants downstream an “attB” attachment site (See Materials and Methods). The combinatorial elements for making the library have been discussed in previous section and in Tables (1-4). The BiTE harboring donor plasmid library was then integrated using an integrase-mediated recombination into the landing-pad of the HEK293_LP^CD19^ cells to obtain a BiTE-expression cell library (HEK293_LP^CD19/BiTE^). A schematic of the donor plasmid integration into the landing pad attachment site is illustrated in Fig. 2b. Following integration, antibiotic selection ensures enrichment of single copy BiTE variant integrated HEK_LP^CD19/BiTE^ cells. Genomic DNA from BiTE integrated cells was selectively PCR amplified with BiTE gene specific primers and PCR amplicons were analyzed by long-read amplicon sequencing (PacBio, SMRT™). We were able to obtain ∼22,300 unique, full length BiTE variants integrated in the mammalian cell library. Additionally, we also confirmed well-distributed representation of design parameters such as scFv clones, linkers and orientations in the integrated BiTE variants (Fig. 2c). We also engineered a single copy Blinatumomab integrated HEK293_LP^CD19^ cell line (HEK293_LP^CD19/Blin^) using the same integrase mediated approach described above with a donor plasmid containing expression cassette of the Blinatumomab BiTE which bears the identical amino acid sequences as the FDA-approved Blinatumomab, including the same C-terminal 6XHis tag. We verified expression of Blinatumomab by flow cytometry using intra-cellular staining of the engineered cells with an anti-his tag antibody (Supplementary Fig. 1c).

### Single-cell-based BsAb functional assay

The core of our single-cell platform is the opto-electro-mechanical module for droplet-based single-cell assay (Fig. 1). This module is capable of highly efficient cell-encapsulation, cell droplet incubation (off-chip in this study), sensitive and efficient detection of intra-droplet live cells using photomultiplier tubes (PMTs), and highly efficient droplet sorting via fluorescence-activated droplet sorting using dielectrophoresis (DEP). Of note, our single cell platform employs innovative multi-point, sequential detections coupled with kinetic droplet indexing, effectively linking functional and phenotypic readouts of individual hits to downstream single-cell genetics (e.g., BsAb sequences), wherein the second-point detection can employ a PMT or serial droplet imaging using a camera to provide spatial resolution. We have demonstrated that individual target and reporter cells can be co-encapsulated with an adequate efficiency of ∼20% (due to Poisson distribution^57^) into uniform stable water-in-oil droplets using a flow-focusing device at a throughput of 1,000s of droplets per second. The droplet size (240 pL used in this study), medium composition and droplet incubation conditions have been tuned to accommodate commonly used cells including Jurkat, HEK293 and primary T cells with ≥90% viability over 9 hours (Supplementary Fig. 2). Intra-droplet fluorescent cells can be detected with ≥95% detection efficiency and the positive droplets can be sorted efficiently even for low-abundance (< 0.1%) clones with a sorting throughput of 100s-1,000 Hz. For example, Supplementary Fig. 3 showed 60-85% sorting efficiency for a starting pool of droplets with 10% positive droplets containing ≥1 fluorescent cells. The sorted droplets can be individually dispensed into microtiter plates for downstream molecular analyses and validations.

**Figure 3.**
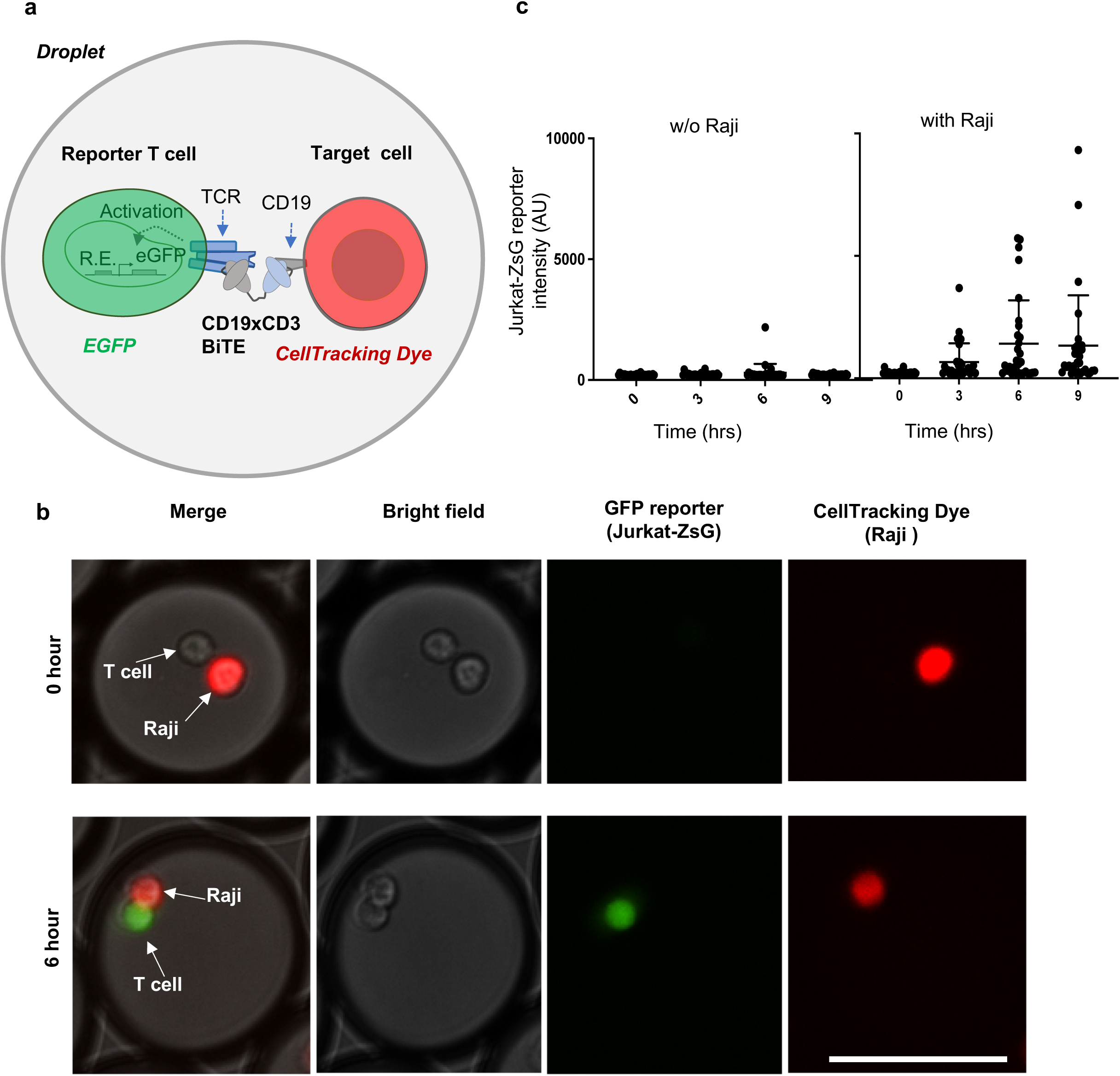
BiTE mediated activation of Jurkat-ZsG reporter cells in droplets. **(a)** Illustrative mechanism of CD19xCD3 BiTE mediated T cell activation in the droplet assay. R.E-responsive element. **(b)** Microscopic images of activated Jurkat-ZsG reporter cells (expressing a GFP reporter) in the presence of Raji cell (CD19+) and a research-grade recombinant Blinatumomab in droplets; scale bar, 100µm. Green, ZsGreen reporter signal; Red, a CellTracking dye pre-labelled for Raji.**(c)** ZsGreen signal intensity change in Jurkat-ZsG cells added with 100 ng/ml Blinatumomab, without(*left*) or with(*right*) over 0-9 hr time-points post incubation. ZsGreen intensity was quantified for *n* = 30 randomly chosen droplets (obtained from one encapsulation experiment) at each timepoint. Data for each timepoint is presented as scatterplot with *mean* + *SD* error bars.

A key innovation of our single cell platform is the ability to directly screen for functionalities that confer therapeutic modes of action that circumvents the need to identify binders first and then evaluate their individual functions as required in conventional assays. Specifically, to screen for functional BiTEs based on T cell activation, we employed a T-cell reporter system (“Jurkat-ZsG”) derived from Jurkat cell (E6.1) that expresses ZsGreen fluorescent protein downstream of TCR activation pathway driven by a promoter with 6x nuclear factor of activated T-cell (NFAT) transcriptionally responsive elements (TREs)^60^ to interrogate BiTE mediated T-cell/tumor cell interaction and T-cell activation (Fig. 3a). We first functionally validated Jurkat-ZsG in a coculture assay with CD19+ Raji cell in bulk using a recombinant BiTE with a sequence identical to that of Blinatumomab and conditioned media from our engineered HEK293_LP^CD19/Blin^ (Supplementary Fig. 4a). Using this bulk assay, we further validated our HEK293_LP^CD19/BiTE^ library secreting functional CD19xCD3 BiTE clones using purified supernatant from HEK293_LP^CD19/BiTE^ cells (Supplementary Fig. 4b). T cell reporter activation mediated by Blinatumomab-BiTE exhibited high activation efficiency (40%) with a false positive rate of 1% and fast kinetics (signal detectable in 3 hours and peaking in 6-9 hours) in droplets (Fig. 3c). These data support that the T cell/target cell interaction mediated by BiTEs can be functionally and efficiently assayed in a compartmentalized droplet format at the single cell level. Similarly, we have developed and characterized a second Jurkat reporter system that expresses a far-red fluorescent protein E2-Crimson (“Jurkat-E2C”) which was used to improve BiTE screening fidelity and reduce false positives. The reporter signal threshold for sorting is typically set to be (µ^negative^ +4σ), where µ^negative^ and σ respectively refer to the mean signal intensity and the standard deviation in unstimulated reporter cells. Of note, the Jurkat reporter cells express very little or no Perforin (a key apoptotic factor) upon T cell activation, and therefore they do not kill the target cells which can hence be retrieved for downstream analyses^61^.

### Assessment of analytical performance of the single cell BsAb discovery platform

We next characterized the throughput, screening efficiency, reproducibility, and false positive rate of our single cell BsAb discovery platform using a spiked library prepared with Blinatumomab expressing HEK293_LP^CD19/Blin^. Briefly, HEK293_LP^CD19/Blin^ was spiked into a pool of parental HEK293_LP^CD19^ cells at different ratios ranging from 0.003% to 1%. Each spiked cell sample is co-encapsulated with the Jurkat-ZsG reporter cells to generate assay droplets using a droplet generation module, where (a) the average number of HEK293 cells per droplet is about 0.3, ensuring that ≥96% droplets contain either zero or one library cell, and (b) ∼78% droplets each have at least one reporter cell, with a theoretically maximal positive-droplet rate of about 31%. Upon six-hour incubation, the assay droplets are subject to detection and sorting with the single cell platform’s core module. Recovered HEK293^CD19/Blin^ cells are verified by single-cell PCR (success rate 75∼80%), followed by Sanger sequencing. Overall, we found that up to 1.5 million cells can be effectively screened per run in a single day. Dispensed droplets can be directly processed or stored at -80°C for batch processing at later time for downstream molecular biology and functional validation.

We observed that, with the HEK293_LP^CD19/Blin^-spiked cell samples, in a total number of 40 experimental runs over different spiking ratios ranging from 0.003% to 1%, our platform can consistently recover up to about 25-30% of all copies of this clone within each of the individual samples, given the theoretical upper limit of recovery rate (p) of about 31% (Fig. 4a). With spiked samples of varying HEK293_LP^CD19/Blin^ abundance (as low as 0.003%), we consistently recovered ≥1 copies of the clone from every run. Through multiple regression modeling (linear, and 4- or 5-parameter logistical), a linear model was selected, which has a significant R^2^ (0.34) with a p-value of 0.0002 and fulfills the statistical assumptions (*e*.*g*., stable residual variance; independence and normality of the residuals). The prediction model with a 90% confidence interval is shown (Fig. 4b). With this prediction model, the screening efficiency (P), defined as the probability of isolating at least one copy of a positive clone, is given by a binomial probability: P = 1-(1-p)^N^ (Fig. 4c), where the Bernoulli probability “p” refers to the recovery rate. The binomial model predicts that our platform is well capable of isolating ≥1 copy of any rare functional clone (as low as 80 copies in one million cells, or 0.008%) with a 95% confidence level.

**Figure 4.**
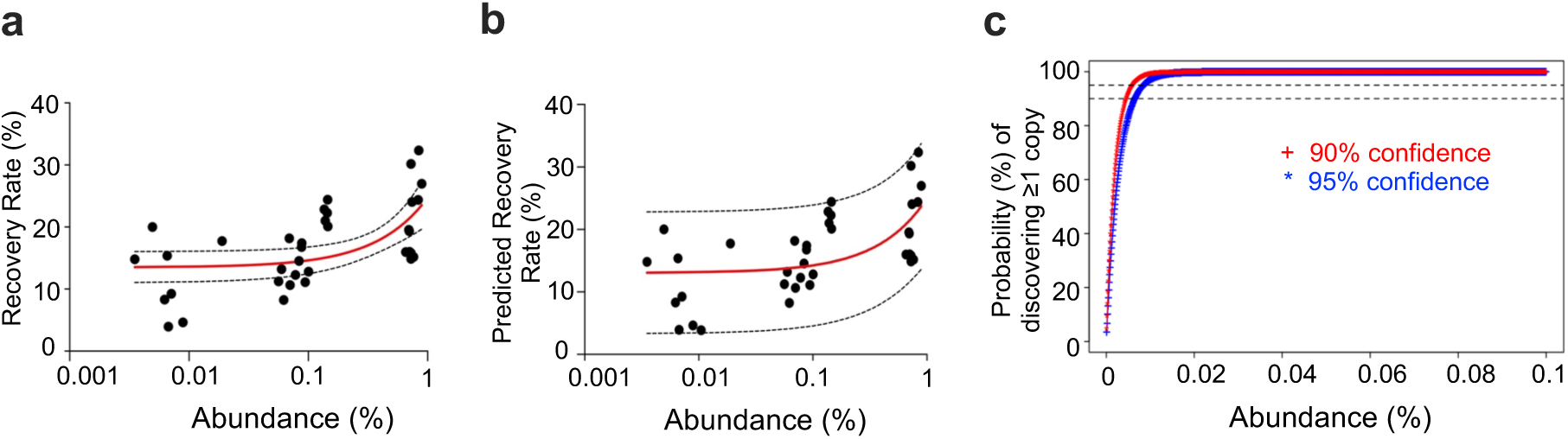
Determination of the screening efficiency for single-cell platform based BsAb functional screening, using HEK293_LP^CD19/Blin^-spiked cell samples. (**a**) Scatterplot showing the observed recovery rate (“p”) versus the abundance of HEK293_LP^CD19/Blin^ spiked in the bulk negative cells (one million HEK_LP^CD19^). Recovery rate is defined as the isolated copy number divided by the total copy number initially spiked into a sample. Each data point (“dot”) is derived from the mix of three analytical replicates of spiked samples. The solid redline is derived from a linear regression with the one standard deviation bounds (dotted lines). (**b**) Predicted recovery rate based on a linear regression model and its 90% confidence interval (R^2^=0.34; p-value=0.0002; statistical assumptions largely fulfilled, e.g., stable residual variance). (**c**) Prediction of screening efficiency (“P”), defined as the probability of discovering ≥1 copy of a positive clone from a cell pool. This model is based on a binomial probability, where the Bernoulli probability “p” refers to the predicted recovery rate in the foregoing regression model, corresponding to a selected confidence lower bound (90% or 95%).

An additional key factor in efficient high throughput screening is to control the false positive rate (FPR), which will impact the downstream workload as well as the operational cost. Reporter cells typically exhibit a certain basal level of inherent, false positive activation. The main source of the false positive activation appears to be non-specific or tonic signaling which may in part attributed to promoter leakiness. For our screening, we typically set a sorting threshold that tolerates 0.1-1% false positives (Supplementary Fig.5a). To reduce FPR, we have employed an orthogonal assay chemistry where both the Jurkat-ZsG and Jurkat-E2C are co-encapsulated with HEK293_LP^CD19/BiTE^, and where only the droplets with dual reporter activation (both ZsGreen and E2-Crimson positive) are sorted. Such orthogonal assay readouts have been found to be highly useful to drastically reduce the false positive rate to <0.01% (Supplementary Fig.5b).

### Discover novel functional BsAbs using single cell discovery platform from a complex library

We next assessed the performance of our single cell BsAb discovery platform for functional BiTE screening from a complex library (HEK293_LP^CD19/BiTE^). As an internal reference, our positive control HEK293_LP^CD19/Blin^ cells were spiked in HEK293_LP^CD19/BiTE^ library cells at 0.1% abundance. All BiTE expressing cells were stained with cell tracking dye (CellTrace™ Violet) (CTV). As described previously, the BiTE-expressing cells are co-encapsulated in droplets with Jurkat reporter cells such that average number of BiTE expressing HEK cells was 0.3 per droplet. For this experiment, we employed a 1:1 mixture of Jurkat-ZsG and Jurkat-E2C in order to reduce false positives. The average number of reporter cells per droplet was about three. Each of “Positive” droplets contains a library cell (CTV^+^, blue peak) and double reporter signals (Jurkat-ZsG and Jurkat-E2C, green and red peaks) after incubating the droplets for about 6 hours (Fig. 5a and Fig. 5b). The ability to interrogate multiple different signals in a multiplex fashion enables us to reduce false positive rate and potentially obtain rare clones of optimal quality. Each individual droplet was initially scanned at the first point of detection (Fig. 1). When a droplet containing double reporter signals (Jurkat-ZsG and Jurkat-E2C) and a HEK293_LP^CD19/BiTE^ library cell signal (CTV) was detected (Fig. 5b), the droplet was pulled into the sorting channel. After that, the sorted droplet moved downstream toward the second point of detection for further confirmation of the fluorescent signals. The second point of detection can employ a PMT or serial droplet imaging using a camera to provide spatial resolution (Fig. 5b). When the sorted droplet passed the thresholds at the second point of detection, a dispensing module was triggered to dispense the single sorted droplet into a PCR tube. The signals of each positive droplet from the first point and second point of detection can be matched with the PCR tube number, so that the signals can be further analyzed after sorting to determine if the sorted droplets are false positive ones. The multiple points of detection and indexing methods can further reduce false positive rate, thereby reducing the overall cost for downstream analysis.

**Figure 5.**
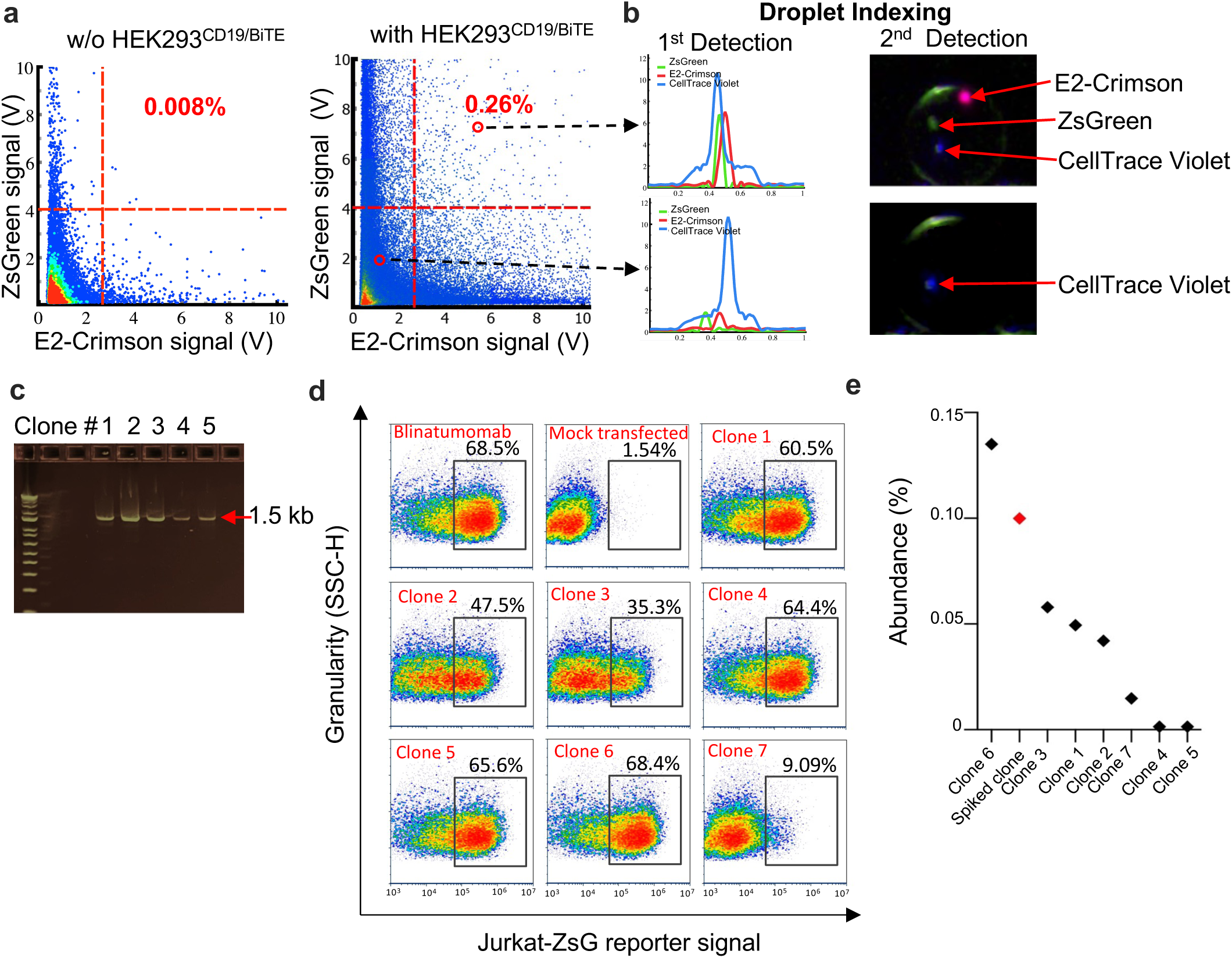
Functional screening of the complex BiTE cell library HEK293_LP^CD19/BiTE^ spiked with 0.1% HEK293_LP^CD19/Blin^ using a double-reporter based orthogonal assay chemistry on the single cell screening platform. **(a)** Representative scatterplots from two independent screening runs showing detected droplet signals after 6-hr incubation. Jurkat-ZsG and Jurakt-E2C reporter cells were co-encapsulated without (*left*) or with (*right*) the BiTE library cells. HEK293_LP^CD19/BiTE^ were pre-labelled with CellTrace™ Violet dye. Gating zones and activation rates are also shown. (**b**) Representative droplet signal (PMT) profiles and images corresponding to a positive (*top)* and negative (*bottom*) cell: HEK293_LP^CD19/BiTE^ cell (CellTrace™ Violet, blue), an activated Jurkat-ZsG cell (ZsGreen, green), and an activated Jurkat-E2C cell (E2-Crimson, red). (**c**) Representative DNA agarose gel image showing single-cell PCR products from a panel of isolated CD19xCD3 BiTE clones. (**d**) Representative flow cytometry plots for bulk functional validation of seven novel BiTE clones (obtained from two screening runs). Jurkat-ZsG reporter cells were co-cultured with CD19+ Raji in the presence of day-3 condition medium harvested from the 1-mL-scale expression from Expi293F™ cells transfected with BiTE gene harboring expression plasmids. After 24-hr incubation, the Jurkat-ZsG reporter cells were profiled by flow cytometry to measure the activation rate. Blinatumomab-BiTE was also transfected and expressed in parallel as a positive control. A total of 265 BiTE clones were tested for functionality using Jurkat-ZsG reporter activation assay(only seven clones are represented here). (**e**) Estimated abundance of select sequence-verified novel BiTEs. Note that Clones 4 and 5 are extremely rare (∼0.001% abundance). The abundance estimation is based on individual clones’ respective counts in the NGS sequences, assuming that these clones are proportionally amplified during the NGS library preparation.

We screened a total of 1.5 million BiTE expressing library cells in two independent runs and dispensed sorted positive droplets each into individual PCR tubes. Of note, the single cell platform consistently recovered the HEK293_LP^CD19/Blin^ (positive control) in each run where we spiked the cells at different ratio among HEK293_LP^CD19^ cells. Fig. 5a shows a representative screening experiment where 0.26% positive hits are detected with a low FPR of 0.008%. Fig.5b shows representative PMT profiles and images corresponding to a positive and negative droplet, respectively, using our tri-color assay (CellTrace™ Violet+ ZsG+ E2C+). We then performed downstream functional characterization of the sorted candidates. For this, we first subcloned BiTE gene fragments obtained by gene specific PCR of cDNA synthesized from each sorted well, into expression plasmids (See Materials and Methods, Supplementary Fig. S6). Fig. 5c shows representative agarose gel image of PCR amplified product from cDNA of representative sorted droplets. We then expressed BiTE clones in 1-mL scale Expi293F culture in a 96-well plate format. 265 clones successfully recovered clones following single-cell PCR and expression were then functionally validated using our reporter cell assay in a conventional bulk format in the presence of Jurkat-ZsG cells and Raji cells (a representative subset of data are shown in Fig. 5d). Using this assay, we confirmed that among the tested 265 clones, 213 clones were true positives, which were then subject to sequencing and further functional characterization including primary T cell activation and tumor cell killing. In addition to the successful isolation of the Blinatumomab analog BiTE (an internal positive control) from the complex library samples, 98 unique CD3xCD19 BiTE sequences were discovered. For example, two clones (Clones 4 and 5) are very rare ones (each at ∼0.001% abundance) (Fig. 5e). Of note, multiple clones including representative Clones 1, 2, and 6 contain a rigid linker (AEAAAKA or AEAAAKEAAAKA). This is quite a surprise, given the conventional wisdom that a flexible linker is generally preferred at the inter-scFv domain interface^62^. Two representative clones, 4G7_VL-VH_-GGGGS-hu38E4.v1_VH-VL_ (Clone 5) and 4G7_VL-VH_-AEAAAKA-L2K_VH-VL_ (Clone 6) with a short flexible and a rigid linker, respectively, were further characterized using a primary T cell and Raji coculture assay (Fig.6a-d) and functionally validated for tumor killing with an efficiency comparable to recombinant Blinatumomab (Fig. 6e). Clone 6 potentiates similar activation and cytotoxic profile from primary T cells as Blinatumomab. Again, Clone 6 possesses a short, rigid linker (AEAAAKA) which to date has not been reported in functional BiTE sequences. These data demonstrate that the high throughput and agnostic nature of our single cell discovery platform enables efficient isolation of rare functional BsAb clones with unique properties, which would otherwise be impossible using conventional low-throughput, trial-and-error methods.

**Figure 6.**
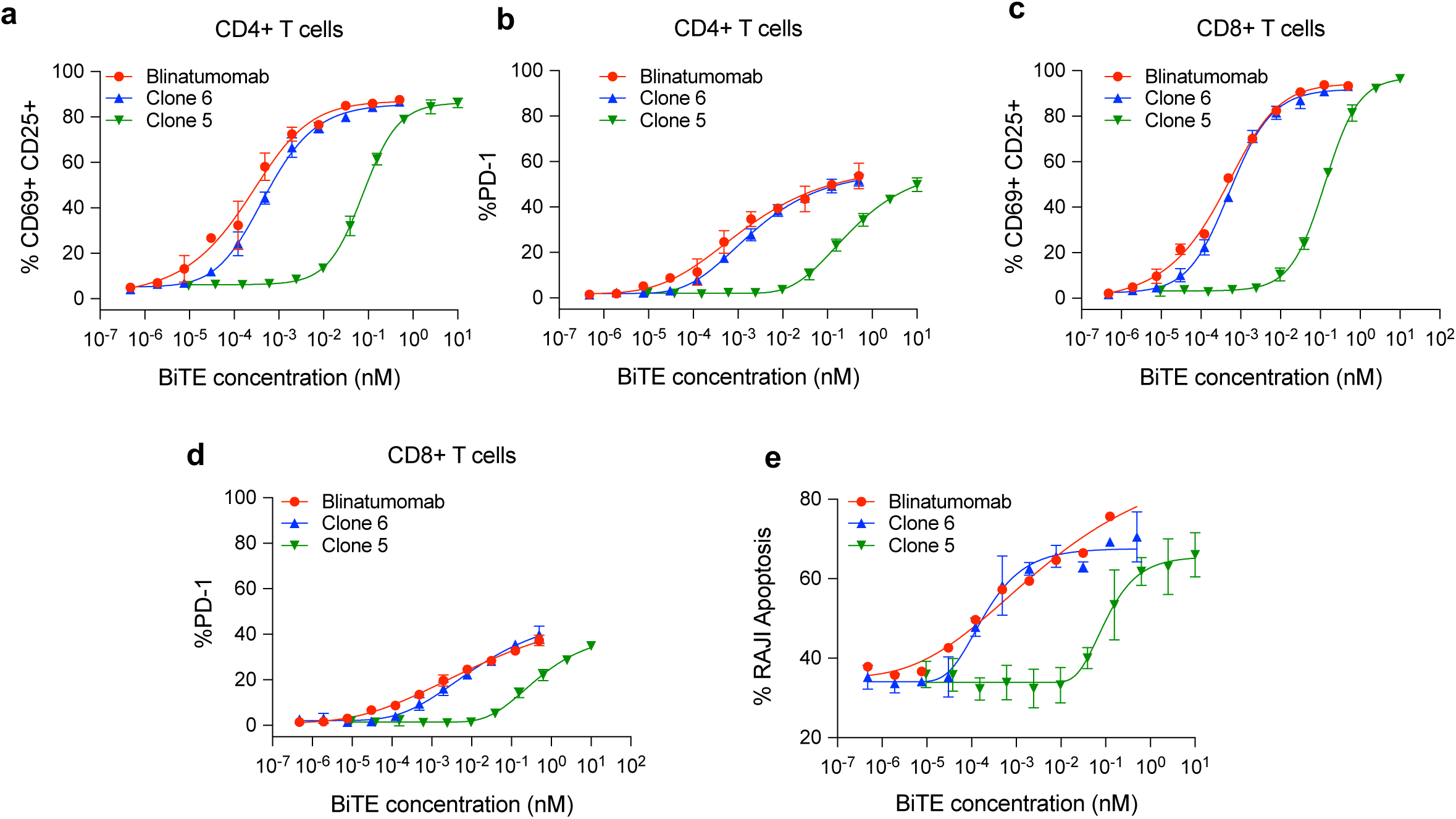
Characterization of activity of purified Clone 5 and Clone 6 with primary T cells. **(a)** Activation profile of CD4+ T cells (assessed by percentage of double positive CD69+CD25+ population) in presence of Raji cells and purified BiTE clones at varying concentrations. **(b)** Percentage PD-1 expressing CD4+ T cells in presence of Raji cells and purified BiTE clones at varying concentrations. **(c)** Activation profile of CD8+ T cells (assessed by percentage of double positive CD69+CD25+ population) in presence of Raji cells and purified BiTE clones at varying concentrations. **(d)** Percentage PD-1 expressing CD8+ T cells in presence of Raji cells and purified BiTE clones at varying concentrations. **(e)** Apoptosis of Raji cells in presence of pan T cells (E:T =10:1), isolated from donor PBMCs and purified BiTE clones at different concentrations assessed by Annexin V/7AAD staining of Raji cells. Data are presented as XY plots and each data point is obtained as mean of *n* = 3 replicates for pan T cells isolated from *n* = 1 donor PBMCs. *mean* + *SD* error bars are plotted for each data point.

### Towards design principles for BsAbs

To elucidate distribution of design variables (which were well distributed in the initial library, Fig. 2c) in the sorted clones that enabled functionality of BiTE variants, we sequenced functional BiTE clones based on the bulk reporter assay using direct bacterial colony sequencing with BiTE gene specific primers and discovered 98 unique sequences (plus the spiked Blinatumomab clone). Interestingly, only two relative orientations of CD19 and CD3 scFvs:

(CD19_VL-VH_–CD3_VH-VL_ and CD19_VH-VL_–CD3_VH-VL_) were represented in the sorted functional candidates, indicating a specific orientational preference for activity (Fig. 7a). Of note, the clinically approved Blinatumomab employs the (CD19_VL-VH_–CD3_VH-VL_) orientation. It is possible that other VH-VL relative orientations including CD19_VL-VH_ -CD3 _VL-VH_ and CD19_VH-VL_-CD3 _VL-VH_ are structurally unfavorable for binding and/or expression^63,64^. Additionally, we also note the underrepresentation (5-6%) of h38E4.v1 CD3 scFv and FMC63 CD19 scFv in the sorted functional library (Fig. 7a) despite their significant representation in the original library. In particular, we do not see any functional BiTE comprising of a h38E4.v1 CD3scFv and FMC63 CD19 scFv in our sorted functional library, irrespective of orientation or linker type (Supplementary Table 1). The underrepresentation of FMC63 CD19 scFv is particularly surprising in that it is the basis for approved CAR-T therapies^47,60,61^, implying that BiTE and CAR-T employ implying that BiTE and CAR-T employ different MOA even with the same scFv potentially due to differences in expression or folding between soluble protein form in BiTE and cell surface protein form in CAR. Furthermore, the association of underrepresentation of FMC63 CD19 with binding epitope or affinity remains to be determined, as 4G7 CD19 scFv, which shares similar binding epitope on CD19 as FMC63 and possesses almost 10-fold lower affinity than FMC63^46,47^, is represented at a level of 36% in the functionally sorted library. This observed differential representation could potentially be due to the protein expression efficiency of different molecular format from host cells (HEK293) in the droplet assay, which will be examined in future work. If true, our single cell discovery platform could not only screen for functionality but also the “developability” (*e*.*g*., efficient expression) of candidate molecules in the same assay, thereby further accelerating down-stream drug development. Furthermore, the four linker types used in the library are well distributed in the sorted clones. Again, rigid linkers are equally functional, contrary to previously thought. We also note that functional screening using the single cell discovery platform can identify key conserved residues in the CDRH3 domain necessary to maintain BsAb functions (Fig. 7b). For instance, we identify residues such as Y10 and Y11 on B43 CDRH3 which are not permissive to mutations and critical for functionality of the BiTE (Fig. 7b). Collectively, these data suggest that our single cell discovery pipeline that interrogates multiple variables in a combinatory, high throughput fashion can, for the first time, inform design principles of BsAbs and reveal the relationship between BsAb sequences at single-residue resolution and resulting function.

**Figure 7.**
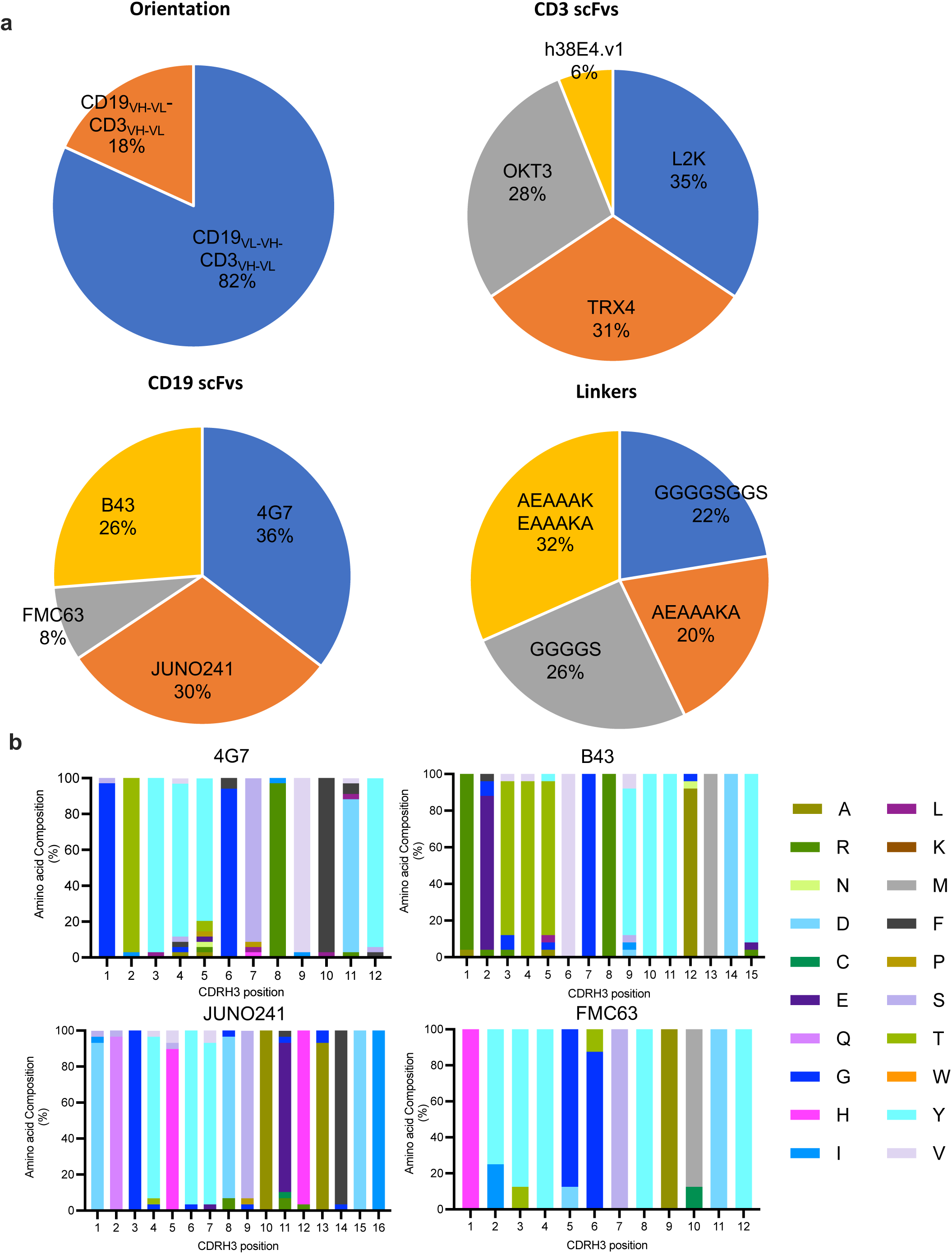
Distribution of design parameters in the functionally active BiTE clones sorted using the single cell discovery platform. **(a)** Distribution of relative orientation of CD19 and CD3 scFvs (*top left*), Distribution of CD3 scFvs (*top right*), Distribution of CD19 scFvs (*bottom left*) and linkers (*bottom right*) in the functionally active sorted library. Sequence of 99 functional BiTE clones(98 unique clones+1 Blinatumomab sequence) elucidated by sanger sequencing were used for analyzing distribution of design parameters. **(b)** CDRH3 composition of sorted BiTEs having a 4G7 (*top left*), B43(*top right*), JUNO241(*bottom left*) or FMC63(*bottom right*) aCD19 scFv. CDRH3 compositions for each CD19 scFv type were analyzed from *n=34* 4G7 containing BiTEs, *n=25* B43 containing BiTEs, *n=29* JUNO241 containing BiTEs and *n=8* FMC63 containing BiTEs. Three clones were excluded from CDRH3 analysis due to sequence ambiguity in the CDRH3 region.

## Discussion

Immunotherapeutics are currently revolutionizing the treatments of many disorders including cancer. BsAbs represent one of the fastest growing sub-sectors in immunotherapy due to their novel modes of action. However, the inefficiency in the current immunotherapeutic discovery paradigm has limited BsAbs’ clinical availability for a broad range of diseases. The conventional trial-and-error approach, in which BsAbs are empirically designed and then must be evaluated individually for their functionality, is generally biased, tedious, time-consuming, and expensive. The single cell BsAb discovery pipeline reported here, differs from conventional microtiter-plate based methods in its working principles by employing a highly parallel, high throughput, unbiased, single cell-based screening which directly interrogates therapeutically relevant functionalities of the candidate molecules. Our platform can accelerate immunotherapeutic pipeline development to potentially make more and better BsAbs quickly available and affordable to patients, enabled by the following superior features compared to conventional methods: (a) Adaptability and scalability: our platform is agnostic, meaning independent BsAb formats can be directly screened for functionality without the need of prior knowledge of BsAb/antigen binding characteristics. (b) Throughput and screening efficiency: our platform can accommodate millions of individual BsAb clones per run, compared to 10s–100s of clones (manual microtiter-plate handling methods) or 1,000s-10,000s of clones (robotics-liquid handling systems) in conventional methods (*i*.*e*., 100– 1,000x increase in throughput). A higher throughput means that one can start with a much bigger library with higher diversity of variants, which therefore increases the success rate in obtaining desired clones, including low-abundance high-quality clones from a large heterogeneous pool. (c) Rapid turnaround: Excluding transitions (*e*.*g*., outsourced NGS services), the estimated turnaround for major processes in a BsAb discovery campaign will comprise: (i) library preparation: 5-6 weeks (wks); and in parallel, reporter cell preparation/validation: 3-5 wks; (ii) 1-2x screening runs: 1-2 days; (iii) downstream validation for ≤96 hits: 3-4 wks. The overall process typically takes 8-10 wks to interrogate 10,000s-100,000s of BsAb variants, compared to ≥10-12 wks required for 100s-1,000 BsAbs using a sophisticated liquid-handling system. (d) Reduced cost: our single cell assay is performed in sub-nanoliter droplets and only requires small sample volume with significantly lower cost of reagents and biological samples.

Other droplet microfluidic systems^65-69^, microwell assays^70^, optofluidics^71^ and microcapillary arrays^72^ have been applied previously for antibody discovery but they either operate with binding-based assays with a moderate throughput or otherwise focused on other areas of discovery not directly applicable to BsAb. A recent study employed single-cell-based functional screening but required multiple rounds of enrichment^73^, whereas innovations in the single cell BsAb discovery platform reported here, including single variant library construction via modular, site-directed landing-pad system, multiplexed orthogonal assay chemistry, and multi-point detection and droplet-indexing strategy enable us to minimize false positives and interrogate large libraries at a much greater depth and lower abundance (0.001%). We have demonstrated that our platform can discover extremely rare clones with unique properties that would otherwise be untractable using conventional low-throughput, biased, and trial-and-error methods.

Structural and compositional parameters in BsAb design may each affect the BsAb function in a delicate yet integral manner. Since our single cell discovery platform can simultaneously interrogate multiple design variables in a combinatorial, high throughput fashion, we can, for the first time, begin to decipher the design principles of BsAbs and obtain in-depth understanding of the inter-relationships among BsAb sequence, structure, and function. Gaining relevant insight into these inter-relationships would enable us to transform development of BsAbs and potentially other immunotherapeutics from a trial-and-error process to a rational engineering practice. We plan to further characterize the novel CD19xCD3 BiTEs discovered from our complex library screening including both in vitro cytotoxicity and in vivo efficacy, which can potentially lead to therapeutic candidates to treat cancer. We will further interrogate the role and mechanism of rigid linkers, preferred VL/VH orientations, scFv’s binding affinity and epitope, conserved residues, and expression efficiency in BiTE functionality. Additional single-cell-based functional assays may further enhance the usefulness of our platform technology, which include immune cell effector function assays relevant to cytokine secretion, metabolism, and/or tumor killing. Such multi-faceted evaluation can facilitate the development of BsAbs that exhibit optimal therapeutic functions for clinical translation. Our single cell platform-based BsAb approach represents a novel route to broadly targeting highly-sought-after yet complex and challenging targets such as a multi-subunit cell-surface receptor.

## Methods

### Preparation of plasmid library of BiTE variants

In order to generate a DNA library pool of BiTE variants in an “attB” containing donor plasmid, we first prepared a plasmid DNA sub library of orientationally switched CD19 and CD3 scFvs (Supplementary Table 2, Table 1). Each sub library candidate was normalized in terminal six residues of the N-terminal and C-terminal portions of both VH and VL domains (Supplementary Table 2). Next, using defined primer pools designed with “NNW” mutations at each CDRH3 position of the CD19 scFvs, we prepared CDRH3 variants of each sub library candidate in the CD19 scFv sub library pool (CD19-NNW-sublibary). Primer sequences encoding for NNW mutations in each residue of CDRH3 of CD19 scFvs are shown in Supplementary Table 3. We next prepared a base plasmid vector which had a multiple cloning site (MCS) consisting of EcoR1 and Kpn1 restriction sites sandwiched between a signal sequence (5’-ATGGGCTGGTCCTGCATCATCCTGTTTCTGGTGGCCACAGCCACAGGCGCCTATGCT-3’) and histag-2A sequence(5’-CACCACCACCATCACCATGCCACGAACTTCTCTCTGTTAAAGCAAGCAGGAGACGTGGAAGAAAACCCCG `GTCCT-3’`). The base plasmid vector additionally had a HindIII restriction enzyme recognition site upstream of the signal sequence and a BamH1 recognition site downstream of histag-2A sequence, to allow for extraction of full BiTE expression cassette (signal sequence-BiTE gene-histag-2A). Using a panel of 16 ssOligos (linker oligos) to encode for the four-member linker library (Supplementary Table 4, Table 3) with appropriate annealing ends to accommodate all the four possible relative orientations of CD19 and CD3 scFvs and four additional oligos annealing to the normalized N-terminal and C-terminal residues of VH or VL domains, we PCR amplified the CD3 variants from CD3-scFv-sublibrary as well as CD19 variants from the CD19-NNW-scFv sublibrary. Primers used for PCR of CD19 variants from CD19-NNW-scFv sub library and PCR of CD3 variants from CD3-scFv-sublibrary are shown in Supplementary table 4. After gel purification and quantification, the PCR products were mixed in equimolar ratios and assembled into linearized base plasmid vector described above, using an in-house Gibson mastermix. Specifically, 100 ng of PCR fragments was assembled with 250 ng of linearized plasmid vector (EcoR1/Kpn1 digested base plasmid vector) in 200 µL Gibson reaction, transformed in 220 µL of DH5α competent cells and recovered in 1mL SOC media for 1 hr without selection. The 1 mL recovered cultures were scaled up 30mL TB media with 100 µg/ml Ampicillin. DNA extracted from the 30mL culture was digested with HindIII-HF (*#R3104S, New England Biolabs*) and BamH1-HF (*#R3136S, New England Biolabs)* and full-length BiTE fragments (∼1.6kb) was extracted using agarose gel extraction. The gel purified full length BiTE fragments were then assembled into a promoterless “attB” donor plasmid. The promoterless “attB” donor plasmid contained the signal Sequence- (EcoR1/Kpn1) MCS-histag-2A-Puromycin cassette downstream of the “attB” attachment site. Linearization of promoterless donor plasmid with EcoR1/Kpn1restriction enzymes allowed for Gibson assembly of gel purified BiTE fragments in-frame with signal sequence and downstream Puromycin sequence. Additionally, we also prepared a Blinatumomab BiTE encoding “attB” donor plasmid vector using B43 encoding CD19 sublibrary plasmid, L2K sequence encoding CD3 plasmid and a ssOligo encoding a short “GGGGS” linker and annealing ends to VH-Cterm and VL-Nterm. Sequence of Blinatumomab used are shown in Supplementary Table 2.

### Engineering HEK293 landing pad cell lines with cell surface expression of CD19

We obtained HEK293 cell lines integrated with a single copy of “attP” attachment site specific to BxB1 recombinase (HEK_LP) from Applied Stem Cell *(#AST-1305, Applied Stem Cell*). We co-transfected our in-house generated Cas9(pCas9) expression plasmid and gRNA plasmid (pGRNA) (specific for integration into Chr 1 locus) with CD19 expression plasmid containing a CAG promoter (pCD19) into the HEK293_LP cell lines. The mass ratio of the plasmids was maintained at 0.33 :0.66:1 (pCas9: pGRNA: pCD19). The transfections were done using Fugene® HD transfection reagent using standard manufacturers protocol.Specifically, 2 µg total DNA was transfected into 1×10^6^ million HEK293_LP cells. Cells were selected using hygromycin and the stable population was sorted on our single cell platform for CD19 expression levels by staining transfected cells with AlexaFluor® 647 anti-CD19 antibody (*#302222, Biolegend*). Cells were sorted for different expression levels of CD19 and clonal cell lines with expression levels comparable to that of standard CD19 positive cell lines such as Raji, NALM-6 and SUDH were chosen for downstream experiments. These cells are referred to HEK293_LP^CD19^.

### Integration of BiTE library into HEK293_LP^CD19^ cells

3×10^6^ HEK293_LP^CD19^ cells were transfected with 60 µg BxB1 recombinase expression plasmid *(#AST 3201, Applied Stem cell*) with BiTE gene integrated “attB” donor plasmid library described above in the ratio 1:3. The transfection was carried out with using Xfect transfection reagent *(#631318, Takara Bio)* as per manufacturer’s protocol. After transfection, the BiTE integrated cells were enriched with selection with 1 µg/ml Puromycin for 48 hrs and was used for downstream droplet sorting and screening or bulk assay.

### Integration of Blinatumomab into HEK293_LP^CD19^

3×10^5^ HEK293_LP^CD19^ cells were transfected with 2.5 µg BxB1 recombinase expression plasmid *(#AST 3201, Applied Stem cell*) with Blinatumomab gene integrated “attB” donor plasmid in the ratio 1:3. The transfection was carried out with using Xfect transfection reagent *(#631318, Takara Bio)* as per manufacturer’s protocol. After transfection, the Blinatumomab integrated cells were enriched with selection with 1 µg/ml Puromycin for 48 hrs and was used for downstream droplet sorting and screening or bulk assay.

### Analysis of diversity of BiTE variants integrated in HEK293_LP^CD19^ cells

Genomic DNA was extracted from 3×10^6^ HEK293_LP^CD19/BiTE^ cells using Quick-DNA midiprep kit *(#D4075, Zymo Research)*. Gene specific primers annealing to the signal sequence (5’-CTGGTCCTGCATCATCCTGTTTC-3’) and P2A sequence (5’-GTTAAAGCAAGCAGGAGACGTGG-3’) were used to amplify BiTE variants from the genomic DNA. The amplification was carried out with 500 ng of template genomic DNA (corresponding to a gene copy number of 145,000) with an annealing temperature of 69 °C and extension time of 1 min 30 sec, using NEB Q5 High-Fidelity DNA Polymerase *(#MO491, New England Biolabs Inc*.). The genomic region containing integrated BiTE sequences amplified by PCR were gel purified and analyzed by long read amplicon sequencing (PacBio SMRT®). The circular consensus reads (∼3 million CCS reads) from long read amplicon sequencing were parsed for gene sequences containing both N-terminal signal sequence and C-terminal his-tag sandwiching the BiTE region using bioinformatics services provided by vendor FornaxBio (*Worcester, MA*). These gene sequences were translated and analyzed for unique and duplicated sequences. The unique BiTE sequences were additionally sorted by linker type, VH-VL orientation of CD19 and CD3 scFvs and representation of scFv candidates. The sequence analysis was carried out using scripts generated in R using standard library packages *(stringr, Biostrings)*.

### Droplet encapsulation of Jurkat reporter cells and HEK293_LP^CD19/BiTE^ cells and sorting

For droplet encapsulation, we used a standard Poisson distribution 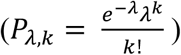 where *λ* is the mean number of cells in a droplet, *k* is the number of cells in the droplets and *P* _*λ,k*_ is the probability that a droplet contains *k* cells, to model the distribution of cells in droplets. Concentration of cells is calculated based on the *λ* needed and average size of droplets (77 µm). We conducted two screening runs on our platform. One sample containing Jurkat-ZsG and Jurkat-E2C reporter cells were mixed in a 1:1 ratio (*λ* = 3) with the other sample containing HEK293_LP^CD19/BiTE^ library cells (*λ* = 0.3) were dispersed into droplets by an immiscible carrier oil using a flow-focusing droplet generator (fabricated in a cleanroom facility at UC Irvine). The carrier oil was Novec HFE7500 fluorinated oil (3M) containing 2% (wt/wt) 008-FluoroSurfactant (Ran Biotechnologies). The HEK_LP^CD19/BiTE^ cells were prestained with CellTrace™ Violet (#C34571, ThermoFisher Scientific) as per manufacturer’s protocol. The flow rates were adjusted to generate droplets with diameter of about 77 µm. Droplets were collected into a capped syringe, and incubated at 37 °C for 6 hrs. After incubation, droplets were reinjected into another microfluidic chip for detecting and sorting on the single cell platform. The single cell platform includes four lasers of wavelength 405 nm, 488 nm, 561 nm and 638 nm at the first point of detection, and two lasers of wavelength 488 nm, and 638 nm at the second point of detection. Droplet fluorescent signals were detected using photomultiplier tubes. The sorting algorithm was programmed by LabVIEW. When a droplet passed the thresholds, a high voltage was triggered to pull the droplet into the sorting channel. The sorted droplet moved to downstream toward the second point of detection for double confirmation of the fluorescent signals. When the sorted droplet passed the thresholds at the second point of detection, a dispensing module was triggered to dispense the single sorted droplet into a PCR tube. The peak profiles of each target droplet detected from PMTs are matched with the PCR tube number containing the target droplet, so that the peak profiles can be manually reviewed to decide to perform downstream analysis, *i*.*e*., single cell PCR, or not. This droplet indexing method further reduces false positive rate and reduce overall cost for downstream analysis. All the droplets that didn’t pass the thresholds were removed to the waste channel.

### Single cell PCR of sorted droplets

cDNA from cells in the sorted droplets was synthesized with total RNA, extracted using QIAGEN RNeasy kit). Specifically, we annealed random hexamer primers to 10 pg of total RNA extracted from single cells in a 20 µL reaction mix containing 1 µL of 50 *μM* random hexamer primers, 1 µL of DNTP mix and 10 pg total RNA. The annealing reaction was conducted at 65 oC for 5 mins. The annealed primer and total RNA was mixed with 4 µL of 5x SSIV buffer (#18090050B, ThermoFisher Scientific), 1 µL 100 mM DTT, 1 µL Ribonuclease inhibitor and 1 µL SuperScript™ IV Reverse Transcriptase (#18090200, ThermoFisher Scientific) and incubated at 23 °C for 10 mins and then at 55 °C for 10 mins. Following this, the reaction was inactivated at 80 °C for 10 mins. Amplification of BiTE gene from the cDNA using BiTE gene specific forward PCR primer annealing to signal sequence(5’-GCTGGTCCTGCATCATCCTGTTTCTG-3’) annealing to the signal sequence and reverse PCR primer annealing to the 2A sequence (5’-CCGGGGTTTTCTTCCACGTCTCC-3’). A second nested PCR was conducted with template from the first PCR to improve product amplification specificity. For this a forward primer annealing to the signal sequence (5’-CTCGGTTCTATCGATTGAATTCCACCATGGGCTGGTCCTGCATCATC-3’) and reverse primer annealing to the histag region (5’-TTGGGCCATGGCGGCCTTAATGGTGATGGTGGTGGTGCTTGATTTC-3’). The primers also added terminal restriction sites (EcoR1 and HindIII) to the PCR product to make it amenable for insertion into expression plasmids. 1 µL linearized expression plasmid was assembled with 3 µL each of PCR amplified product using 4 µL of NEBuilder HiFi DNA Assembly MasterMix *(#E2621L, New England Biolabs)* in a 96-well plate format. The assembled gene products were directly transformed into 30 µL of competent XL1-Blue cells and recovered for 30 mins in 200 µL SOC without selection, in a 96 well plate format. The recovered bacterial cells from each well was inoculated in 2 mL of TB media with 100 *ug*/*ml* Ampicillin for 12-16 hrs at 37 °C. Following overnight incubation, DNA was extracted from the bacterial cells using ZymoPURE plasmid miniprep kit (#*D4212, Zymo Research*) using manufacturer’s protocol. Concentration of each of the extracted DNA was quantified.

### Small scale transient expression of sorted BiTE candidates

The sorted BiTE candidates assembled into expression plasmids, were then transfected into 1mL Expi293F™ cells (*#A14527, ThermoFisher Scientific)* using Expifectamine™ 293 Transfection kit *(#A14524, ThermoFisher Scientific)* as per manufacturers protocol in 96-well 2-mL culture plates. 72 hours after transfection, BiTE supernatant was collected and used for bulk reporter activation assay.

### Bulk activation reporter assay

72-hr BiTE containing supernatant (described above) was mixed with 5×10^4^ Jurkat-ZsG cells and 5×10^4^ Raji cells for 24 hrs in a 96 well V-bottom plate. The cells were then washed with PBS and analyzed for zsGreen fluorescent protein expression. Population of activated Jurkat-ZsG cells were identified based on gating strategy described in Supplementary Fig. 7.

### In-vitro activation and cytotoxicity of select BiTE clones

Pan T cells were isolated from donor PBMC’s (#70025, StemCell Technologies) using EasySep™ human T cell isolation kit (#17951, StemCell Technologies). 10,000 Raji cells were mixed with 100,000 T cells in media containing different concentrations of purified BiTE clones (Clone 5 and Clone 6). After 48hrs, half of the cells were collected and stained with 7AAD (#50-850-582, Fisher Scientific) and Annexin V (#640906, Biolegend) and PE/Cyanine 7 anti-human CD4 antibody (#317414, Biolegend), APC/Cyanine 7 anti-human CD8 antibody (#344714, Biolegend). Population of Annexin V+and 7AAD+ Raji cells obtained by gating out T cells (CD4+/CD8+) (Supplementary Fig. 8). The other half was stained with cocktail containing 7AAD, PE-anti human CD69 antibody (#310906, Biolegend), FITC-anti human CD25 antibody (#356106, Biolegend), APC anti-human CD279 antibody (#329908, Biolegend), PE/Cyanine 7 anti-human CD4 antibody (#317414, Biolegend), APC/Cyanine 7 anti-human CD8 antibody (#344714, Biolegend). Total apoptosis of Raji cells was obtained by gating for 7AAD+ and Annexin V+ population after gating out T cells (CD4+CD8+). Percentage of CD4+CD69+CD25+ T cells, CD4+PD-1+ T cells, CD8+CD69+CD25+ T cells, CD8+PD-1+ T cells was obtained by gating for subpopulation of CD4+ only T cells and CD8+ only T cells from live (7AAD-) T cells (Supplementary Fig. 9). All antibody concentrations were used at concentration recommended by vendors.

### Sequence elucidation of functional BiTE variants

BiTE expression plasmids corresponding to functional BiTE variants were retransformed in XL1-blue competent cells and plated on to LB Agar Ampicillin (100 *μg*/*ml*). Three colonies per BiTE candidate was inoculated for 6 hrs at 37 °C into 500 *μl* of TB Ampicillin (100 *μg*/*ml*) in 96 well 2-mL bacterial culture plates. 100 *μl* of the culture was pipetted into 96-well PCR plate and used for RCA template generation and sanger sequencing. Service for RCA template generation and sanger sequencing was provided by ELIM Biopharm (*Hayward, CA)*. For sanger sequencing we used forward primer (5’-AACATCCACTTTGCCTTTCTCTCC-3’) annealing to signal sequence and reverse primer (5’-GGACAAACCACAACTAGAATGCAG-3’) annealing to histag.

### Purification of novel BiTE clones

Two sorted BiTEs with respectively short rigid and flexible linkers (4G7_VL-VH_-AEAAAKA-L2K_VH-VL_ and 4G7_VL-VH_-GGGGS-hu38E4.v1_VH-VL_ were expressed from a 30 mL transfection scale. The BiTEs were purified from day 3 conditioned medium using the C-terminus 6XHis tag and immobilized metal affinity chromatography (Cytiva, #29051021) on ÄKTAExplorer100 FPLC system (Cytiva). After elution with an Imidazole gradient, the BiTE proteins were buffered exchanged to PBS and concentrated using Amicon Ultra-15 ultracentrifugation 10kDa cutoff tubes (Millipore Sigma, #UFC901096D).

## Author contributions

A.S., Y.S. G.W. and W.Z. designed the study. X.C., P.N.H., J.C., and Q.F. developed the droplet microfluidic system. Y.S., G.W., A.S., J.Z., J.J designed the CD19xCD3 BiTE library. J.J., J.Z. and X.J. generated the DNA library; J.J. and A.S. engineered the mammalian cell library, CD19+ and Blinatumomab+ cells. J.C., A.F., X.C. and A.S. performed the spiking experiments. J.C., A.F., X.C., Q.F. and A.S. performed the library screening experiment. R.L. and J.J. did the downstream analysis (scPCR, DNA recovery and assembly for expression, sequence analysis, small scale expression). J.J. and A.S. ran the functional validations of sorted clones, and J.J. did the sequence analysis. A.F. and R. R. expressed and purified the BiTEs. A.S., X.M. and C.R. did the primary T cell assays for sorted BiTE validation. A.S., J.J., X.C., G.W., W.Z., analyzed the data and wrote the manuscript.

## Acknowledgement

This work is supported by NSF/SBIR grants (#1913404 and #2051931). We acknowledge Dr. Melanie Oaks (UCI Genomics High Throughput facility) for helpful discussions on processing NGS sequencing data.

