## Supplementary Figures for "A high throughput bispecific antibody discovery pipeline"

### co-first authors

$ co-second authors

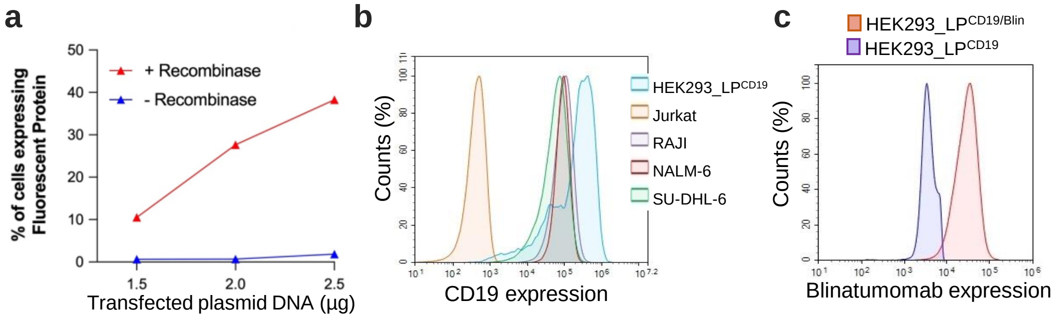


**Supplementary Figure 1**. **Establishment of a HEK293 cell line expressing CD19 (target) and a positive CD19xCD3 BiTE (Blinatumomab).** (**a**) Line graph showing the recombinase-dependent integration efficiency at the landing pad of the HEK293 cells, using a mCherry-encoding donor plasmid as a test material. (**b**) Flow cytometry profiling of a selected clone showing moderate to high CD19 ("HEK293_LP$^{CD19}$"), compared to a few lymphoma lines (RAJI, NALM-6 and SU-DHL-6); Jurkat served as a negative control. (**c**) Flow cytometry validation of HEK293_LP$^{CD19/Blin}$ cells with a CD19xCD3 BiTE (Blinatumomab) that is integrated into the landing pad. The expression of Blinatumomab (6xHis-tagged) was validated using the cell staining with an anti-6xHis antibody.

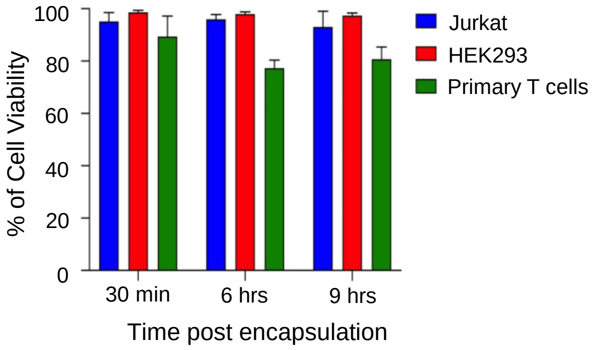


**Supplementary Figure 2.** The viability profile of a few human cell types over a duration that is typical of screening assays conducted on single cell discovery platform. Cells are respectively encapsulated in 250-pL droplets and incubated at 37$^{o}$C under 5% CO2. Droplets are de-emulsified to recover cells in bulk for subsequent 7AAD staining followed by flow cytometry. Jurkat (E6.1), a human leukemia T cell line; HEK293, an immortalized embryonic kidney epithelial cell line; primary T cells: enriched from human PBMCs. Values of mean from $n = 2$ encapsulation experiments are presented as bar graphs with $mean \pm SD$ error bars.

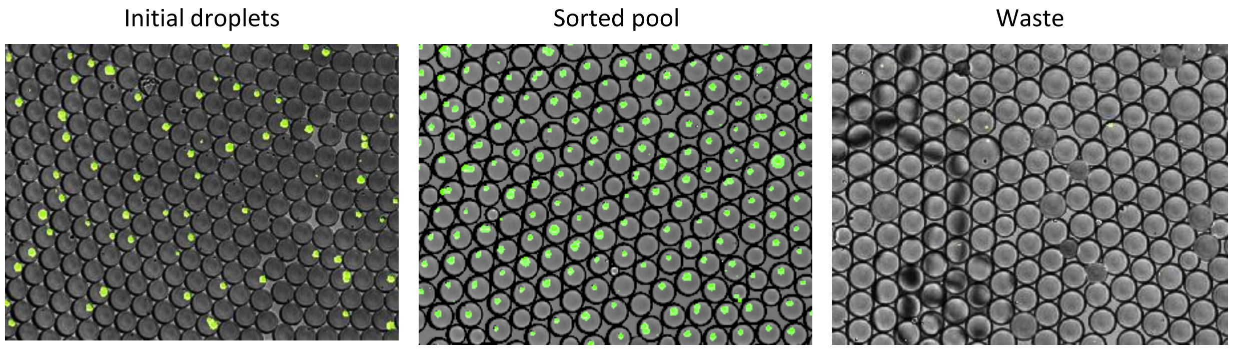


**Supplementary Figure 3. Microscopic images for droplets before sorting** (*left*), sorted pool(*center*), and waste pool(*right*). Green, GFP reporter signal in activated individual reporter T cells (Jurkat) inside droplets (bright field image). Scale bar, 200 µm.

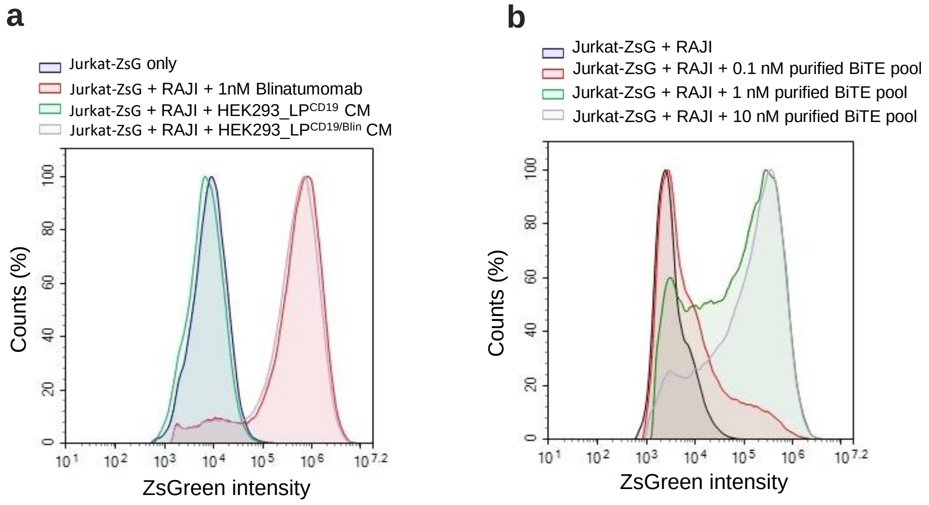


**Supplementary Figure 4. Functional validation of secretion of Blinatumomab and BiTE library by assay of cell culture supernatants. (a)** Blinatumomab containing cell culture supernatants (CM) mixed with a co-culture of Jurkat-ZsG and Raji cells for 24 hrs activated Jurkat-ZsG reporter cells. Media containing 1nM recombinant Blinatumomab is used as positive control. (**b**) Jurkat-ZsG reporter cells were co-cultured with HEK293$^{CD19}$ for 24 hours in the presence of purified BiTE pool (0.1, 1 and 10 nM respectively) from the day-3 condition medium of the library cell culture. Reporter activation was measured using flow cytometry.

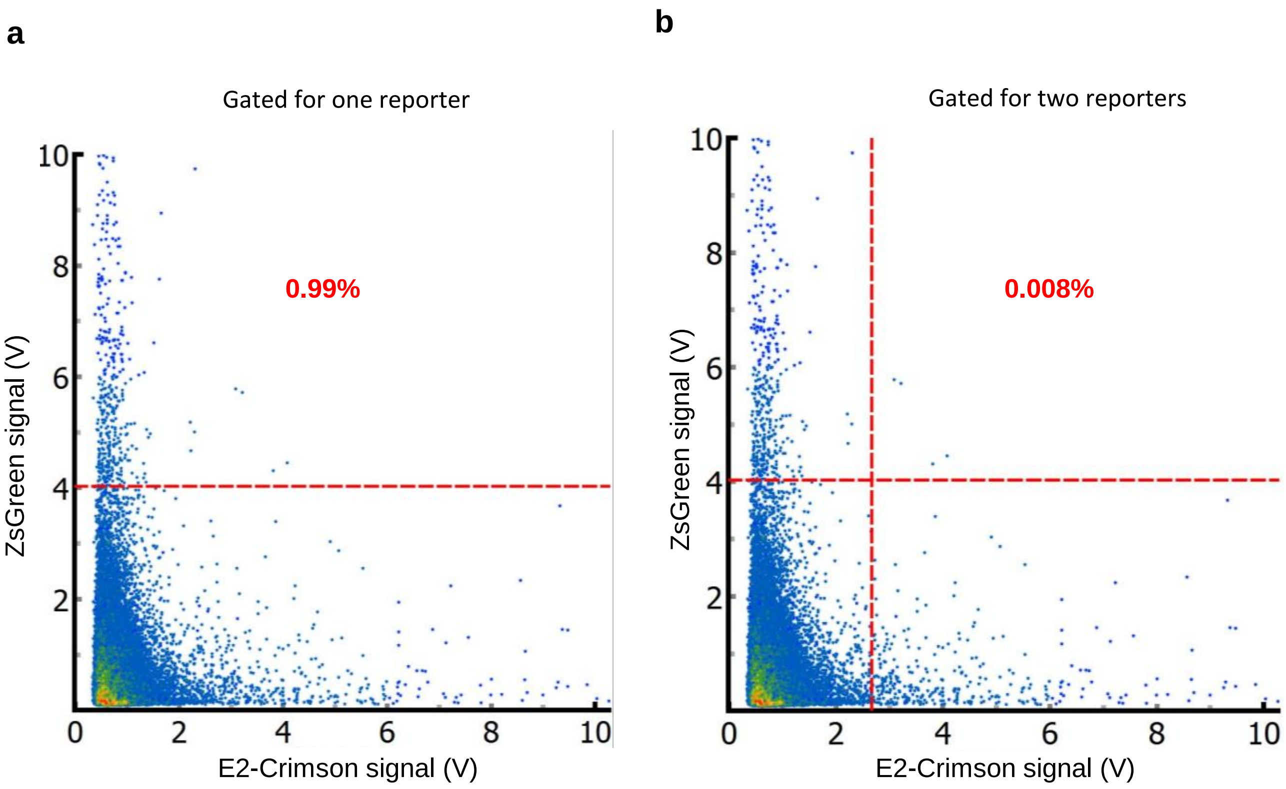


**Supplementary Figure 5. Representation of false positivity rate from detection modules of the single cell discovery platform (a)** False positive rate when only ZsGreen reporter signal is used. (b) False positive rate when both ZsGreen and E2-Crimson signal are used.

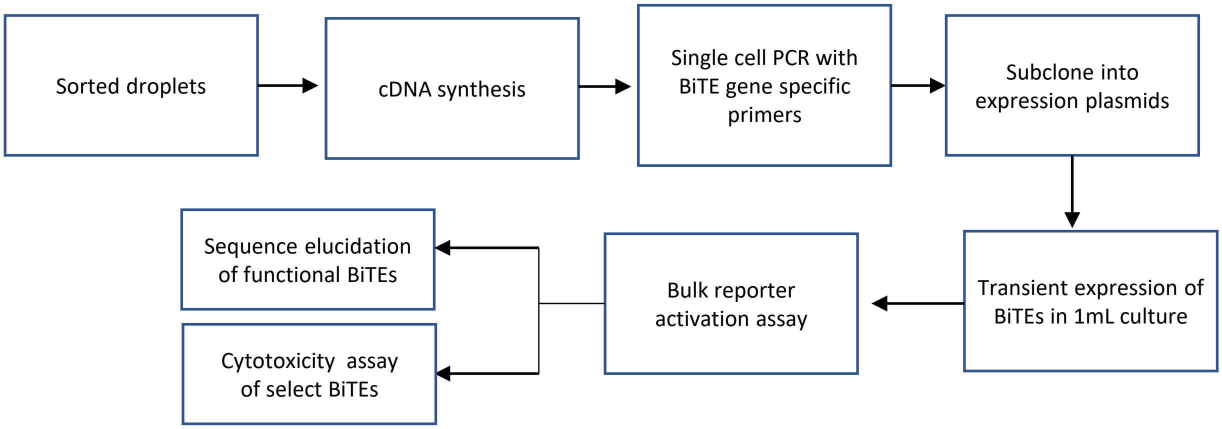


**Supplementary Figure 6. Flowchart illustrating downstream processes after sorting.**

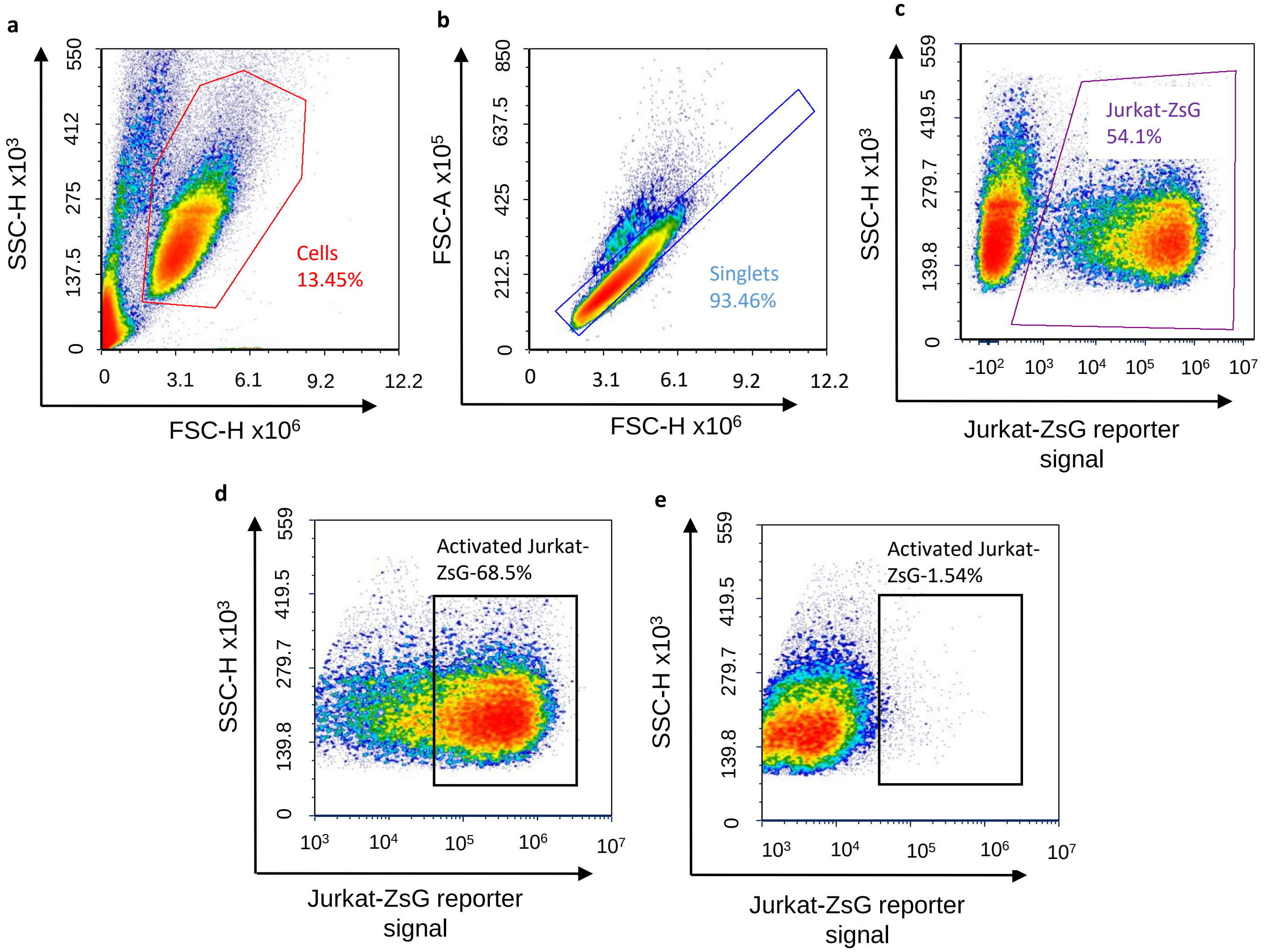


**Supplementary Figure 7. Flow cytometry gating strategy for bulk reporter activation assay**. BiTE clones were expressed in Expi293F$^{TM}$ cells and conditioned media from Day 3 expression was used in a coculture of Raji and Jurkat ZsG cells. 24 hrs after co-culture, ZsGreen protein expression from Jurkat-ZsG cells was analyzed by flow cytometry. Representative gating strategy for identifying population of activated Jurkat-ZsG cells is shown for the case where Raji and Jurkat-ZsG cells were co-incubated with Blinatumomab conditioned medium.**(a)** Cell debri was first excluded based on FSC-H/SSC-H gating. **(b)** *S*inglets were isolated with FSC-H/FSC-A gates*.* **(c)** Raji cells were then excluded based on basal ZsG expression from Jurkat-ZsG reporter cells. **(d)** Percentage of activated reporter cells *(bottom left)* were identified by gating for Jurkat-ZsG cells expressing ZsGreen over basal level. **(e)** A negative control consisting of Jurkat-ZsG and Raji cells co-incubated with media from mock transfected Expi293F$^{TM}$ cell line was used to identify basal Jurkat-ZsG fluorescence.

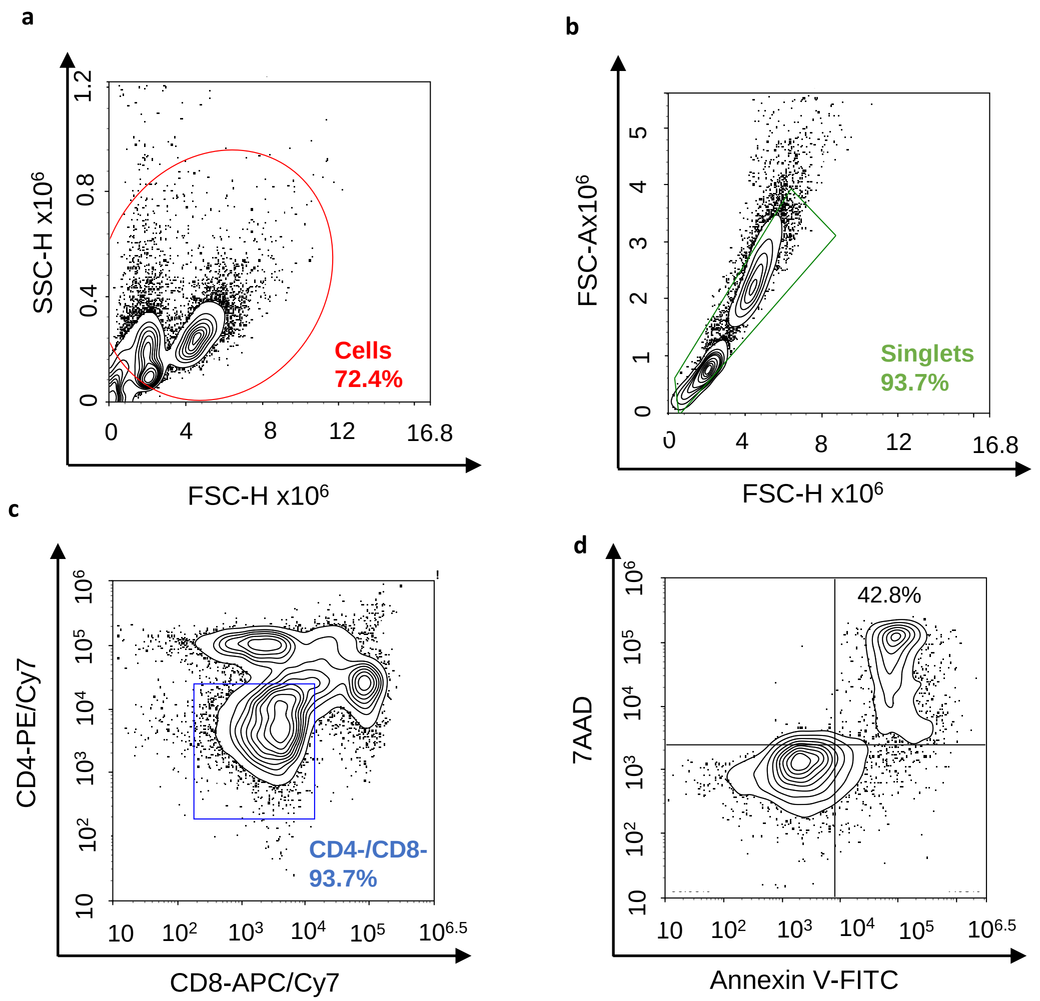


**Supplementary Figure 8**. **Flow cytometry gating strategy for in-vitro cytotoxicity assay**. PanT cells isolated from donor PBMCs were co-incubated with Raji cells in presence of purified BiTE clones at different concentrations. 48 hrs after co-culture, cytotoxicity was assessed by flow cytometry. Gating strategy for the flow cytometric analysis is shown for a representative case where panT cells and Raji cells are co-incubated with recombinant Blinatumomab(used as positive control at 0.125 nM). **(a)** Cell debri was excluded with FSC-H/SSC-H gating. **(b)** Singlets were identified using gates for FSC-A/FSC-H. **(c)** Raji cells were identified as a non-T cell population(double negative CD4-CD8- population). **(d)** Annexin V+ and 7AAD+ cells were identified from Raji cell population *(top right quadrant).*

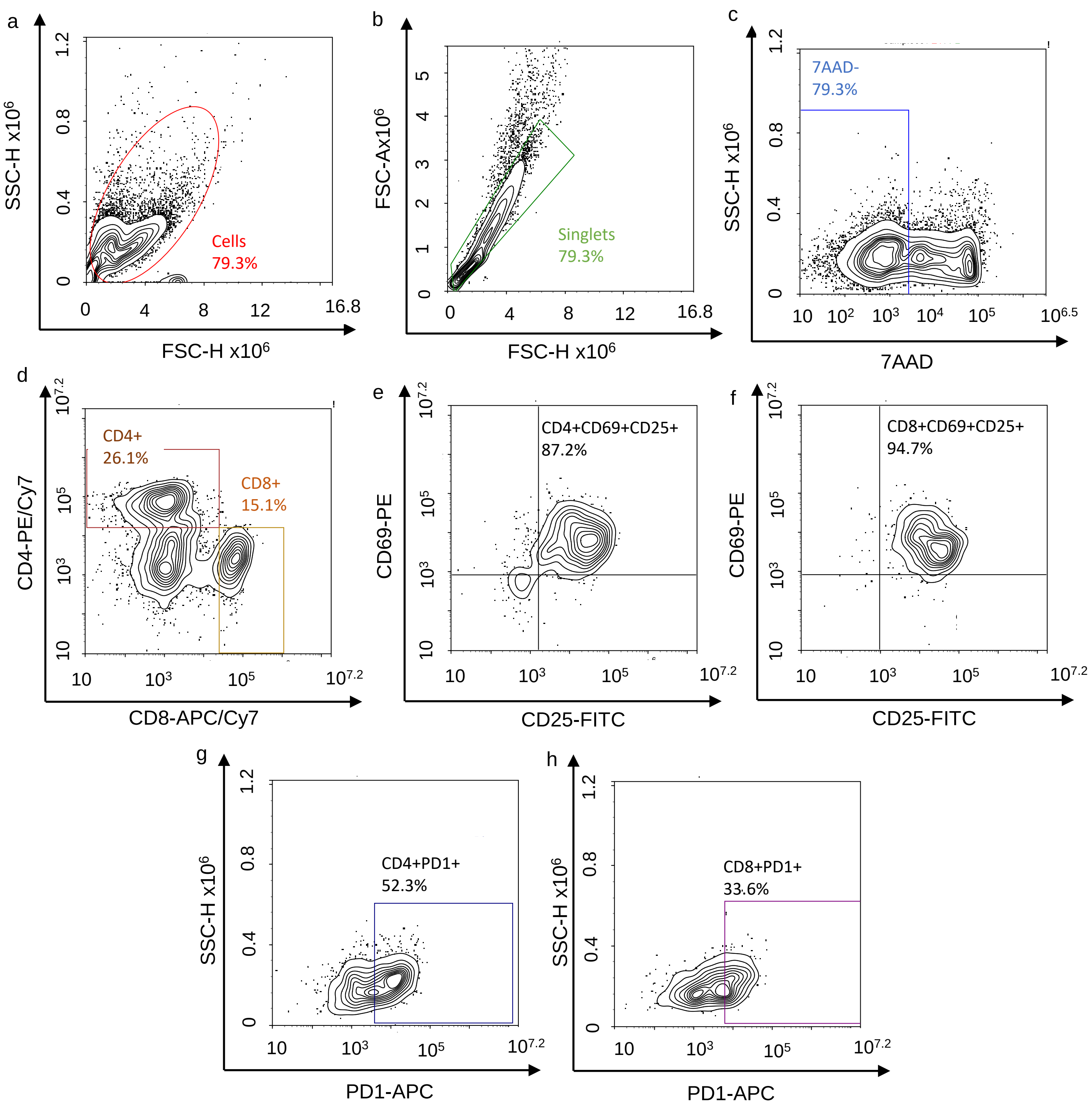


**Supplementary Figure 9**. **Flow cytometry gating strategy for in-vitro cytotoxicity assay**. PanT cells isolated from donor PBMCs were isolated from co-incubated with Raji cells in presence of purified BiTE clones at different concentrations. 48 hrs after co-culture, activation was assessed by flow cytometry. Gating strategy for the flow cytometric analysis is shown for a representative case where panT cells and Raji cells are co-incubated with recombinant Blinatumomab(used as positive control at 0.125 nM). **(a)** Cell debri was excluded with FSC-H/SSC-H gating. **(b)** Singlets were identified using gates for FSC-A/FSC-H. **(c)** 7AAD- population was used to gate for live cells **(d)** CD4+ and CD8+ cells were identified. **(e)** Percentage of CD4+CD69+CD25+ cells were identified by subgating for CD69+ and CD25+ from the CD4+ population**. (f)** Percentage of CD8+CD69+CD25+ cells were identified by subgating for CD69+ and CD25+ from the CD8+ population **(g)** Percentage of CD4+PD1+ population was obtained by subgating for PD1+ from the CD4+ population (h) Percentage of CD8+PD1+ population was obtained by subgating for PD1+ from the CD8+ population.
